# MHC class II-restricted antigen presentation is required to prevent dysfunction of cytotoxic T cells by blood-borne myeloids in brain tumors

**DOI:** 10.1101/2022.06.10.495502

**Authors:** Michael Kilian, Ron Sheinin, Chin Leng Tan, Christopher Krämer, Mirco Friedrich, Ayelet Kaminitz, Khwab Sanghvi, Katharina Lindner, Frederik Cichon, Stefanie Jung, Kristine Jähne, Miriam Ratliff, Robert Prins, Andreas von Deimling, Wolfgang Wick, Asaf Madi, Lukas Bunse, Michael Platten

## Abstract

Cancer immunotherapy critically depends on fitness of cytotoxic and helper T cell responses. Dysfunctional cytotoxic T cell states in the tumor microenvironment (TME) are a major cause of resistance to immunotherapy. Intratumoral myeloid cells, particularly blood-borne myeloids (bbm), are key drivers of T cell dysfunction in the TME. We show here that major histocompatibility complex (MHC) class II (MHCII)-restricted antigen presentation on bbm is essential to control the growth of brain tumors. Loss of MHCII on bbm drives dysfunctional intratumoral tumor-reactive CD8^+^ T cell states through increased chromatin accessibility and expression of Tox, a critical regulator of T cell exhaustion. Mechanistically, MHCII-dependent activation of CD4^+^ T cells restricts myeloid-derived osteopontin that triggers a chronic activation of *nuclear factor of activated T cells* (Nfat)2 in tumor-reactive CD8^+^ T cells. In summary, we provide evidence that MHCII-restricted antigen presentation on bbm is a key mechanism to directly maintain functional cytotoxic T cell states in brain tumors.

**Highlights:**

- MHCII on intratumoral blood-borne myeloid cells is required for sustained anti-glioma T cell response
- Loss of myeloid MHCII drives dysfunction of CD8^+^ T cells by activating TOX
- Tox expression is induced by osteopontin-triggered chronic NFAT2 signaling in tumor-reactive CD8^+^ T cells
- Osteopontin production is restricted upon intratumoral T helper cell activation via MHCII
- MHCII expression on tumor-infiltrating bbm correlates with osteopontin expression and cytotoxic T cell dysfunction in human glioblastoma tissue

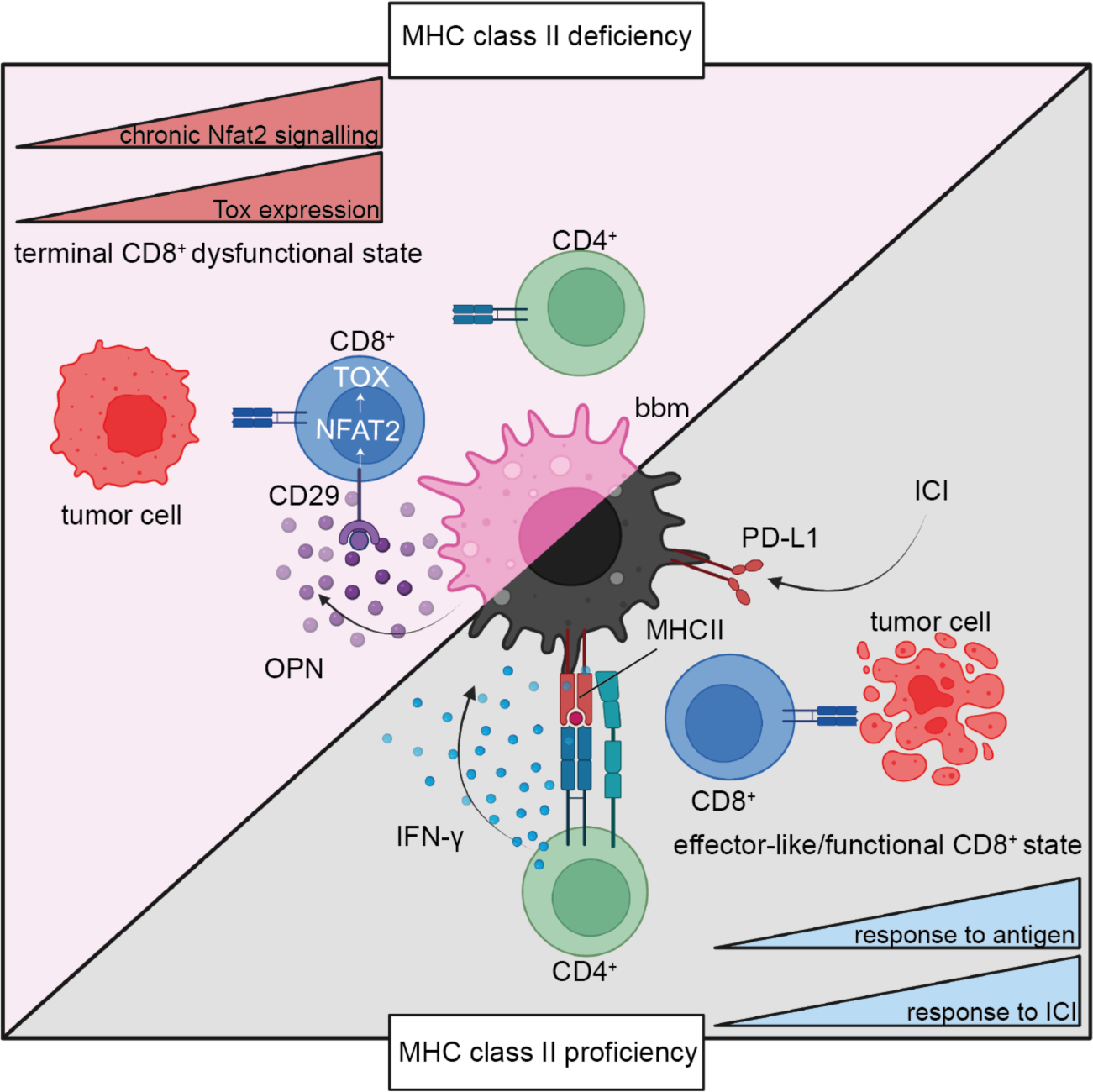

## Main Text

Clinical responses against tumors correlate with cytotoxic T cell infiltration and neoantigen burden (Chen and Mellman, 2017; Schumacher and Schreiber, 2015). Failure of cancer immunosurveillance is thus causally related to insufficient and dysfunctional cytotoxic T cell responses (Thommen and Schumacher, 2018). Intratumoral CD8^+^ T cells display very heterogenous transcriptional and functional phenotypes (Chihara et al., 2018; van der Leun et al., 2020; Li et al., 2019; Singer et al., 2016). Upon antigen-recognition, tumor infiltrating CD8^+^ T cells have been shown to undergo rapid chromatin remodeling, eventually resulting in T cell exhaustion and dysfunction (Pauken et al., 2016; Philip and Schietinger, 2021; Philip et al., 2017; Wherry and Kurachi, 2015; Zheng et al., 2022). Therefore, maintenance and restauration of tumor-reactive T cell fitness remains the key challenge in immune-oncology. Chronically activated T cells upregulate expression of checkpoint receptors, such as PD-1, CTLA-4, TIM-3 and LAG3 (Pardoll, 2012). Immune checkpoint blockade aims at blocking these checkpoint receptors and restoring T cell functionality and has revolutionized cancer therapy (Ribas and Wolchok, 2018; Sharma and Allison, 2015). Response to immune checkpoint inhibition (ICI) is associated with distinct T cell states that are prerequisites for effective immunotherapy (Sade-Feldman et al., 2018; Zhang et al., 2021).

Particularly in central nervous system (CNS) tumors growing in an immune-privileged site, induction of intratumoral T cell responses remain challenging even though robust CD8^+^ or CD4^+^ peripheral T cell responses can be induced (Hilf et al., 2019; Keskin et al., 2019; Platten et al., 2021; Weller et al., 2017) and neoadjuvant treatment of PD-1 checkpoint blockade led to promising responses in some glioblastoma (GBM) patients (Cloughesy et al., 2019; Schalper et al., 2019; Zhao et al., 2019). This is due to a scarcity of neoantigens in GBM cells as well as the distinct immunosuppressive microenvironment in brain tumors. Oncometabolites like R-2-hydroxyglutarate or proteins like osteopontin or Il-10 released in the tumor microenvironment (TME) have been shown to suppress anti-tumor immune responses (Bunse et al., 2018; Friedrich et al., 2021; Nduom et al., 2015; Wei et al., 2019). In experimental gliomas, response to ICI is dependent on both, T helper and cytotoxic T cell responses (Aslan et al., 2020). Similarly, MHC class I- or class II (MHCII)-restricted antitumor vaccines require tumor-reactive CD4^+^ or CD8^+^ T cell responses, respectively, to eradicate solid tumors (Alspach et al., 2019; Kreiter et al., 2015). CD4^+^ T cell responses promote induction of cytotoxic CD8^+^ T cell responses and mediate the formation of a memory CD8^+^ T cell pool required for sustained anti-tumor immune responses (Borst et al., 2018; Laidlaw et al., 2016).

In this study we aimed at determining the impact and mechanistic contribution of intratumoral MHCII-restricted antigen presentation to brain tumor immunosurveillance. We found that intratumoral MHCII is required for sustained cytotoxic CD8^+^ T cell responses and that loss of MHCII drives a distinct transcriptional state in tumor reactive CD8^+^ T cells. Furthermore, we provide evidence that CD4^+^ T cells suppress osteopontin expression in tumor-infiltrating blood borne myeloid (bbm) cells which drives TOX-mediated CD8^+^ T cell dysfunction by stimulating chronic NFAT2 activation.

## Results

### Response to ICI is dependent on MHC class II expression on blood-borne myeloid cells but not brain-resident microglia

Neoadjuvant ICI has recently been demonstrated to improve overall survival in GBM patients (Cloughesy et al., 2019). In experimental gliomas, response to ICI is dependent on CD4^+^ T cells (Aslan et al, 2020). Therefore, we hypothesized that MCHII expression in gliomas impacts response to ICI. By RNA sequencing (RNA-seq) of GBM tissues from patients treated with neoadjuvant aPD-1 (Cloughesy et al., 2019), we defined MHC^high^ and MHC^low^ expressing gliomas and observed a favorable clinical course in patients with MHCII^high^ compared to MHCII^low^ tumors (**Figure 1A**). However, bulk RNA-seq was neither able to investigate spatial nor cellular heterogeneity of MHCII expression. To spatially resolve MHCII expression, we performed in situ RNA hybridization in GBM tissue of treatment-naïve patients. We observed MHCII expression primarily in *Integrin Subunit Alpha M* (ITGAM, CD11b)^+^ myeloid cells and a high inter-patient variability in MHCII expression. (**Figure 1B**). Brain-resident microglia and tumor-infiltrating bbm are not only the most abundant cell types in the brain TME but represent bona fide MHCII antigen presenting cells (Friedrich et al., 2021). To assess cellular MHCII heterogeneity, we first interrogated publicly available human single cell (sc)RNA-seq glioma datasets and found an increased MHCII antigen presentation signature (see STAR methods) on bbm compared to microglia (**Figure 1C, D**, **Figure S1 A**) (Darmanis et al., 2017). Using time-resolved scRNA-seq of murine tumor-infiltrating immune cells from early (d14) and late stage (d26) syngeneic GL261 gliomas we observed constant MHCII expression by murine microglia over time and overall high but decreasing MHCII expression in late stage bbm indicating a dynamic process of MHCII expression **(Figure S1B, C, D)** (Friedrich et al., 2021). To validate that the differential scRNA-seq-retrieved antigen presentation signature has functional implications for T cell activation in our mouse model, we performed *ex vivo* co-culture assays with CD45^high^ CD11b^high^ bbm and CD45^int^ CD11b^high^ microglia and OT-II T cells, recognizing the MHCII model antigen chicken ovalbumin 323-339 (OVA^323-339^) (**Figure S1E, F**). Indeed, using *ex vivo* bbm and microglia comparatively as antigen presenting cells, IFN-γ production in OT-II T cells was increased when recognizing OVA^323-339^ on bbm compared to microglia (**Figure S1G**). Taken together, in gliomas, MHCII expression correlates with response to ICI and is heterogenous on a spatial and cellular level. Particularly on bbm, MHCII expression is high and dynamic.

**Figure 1.**
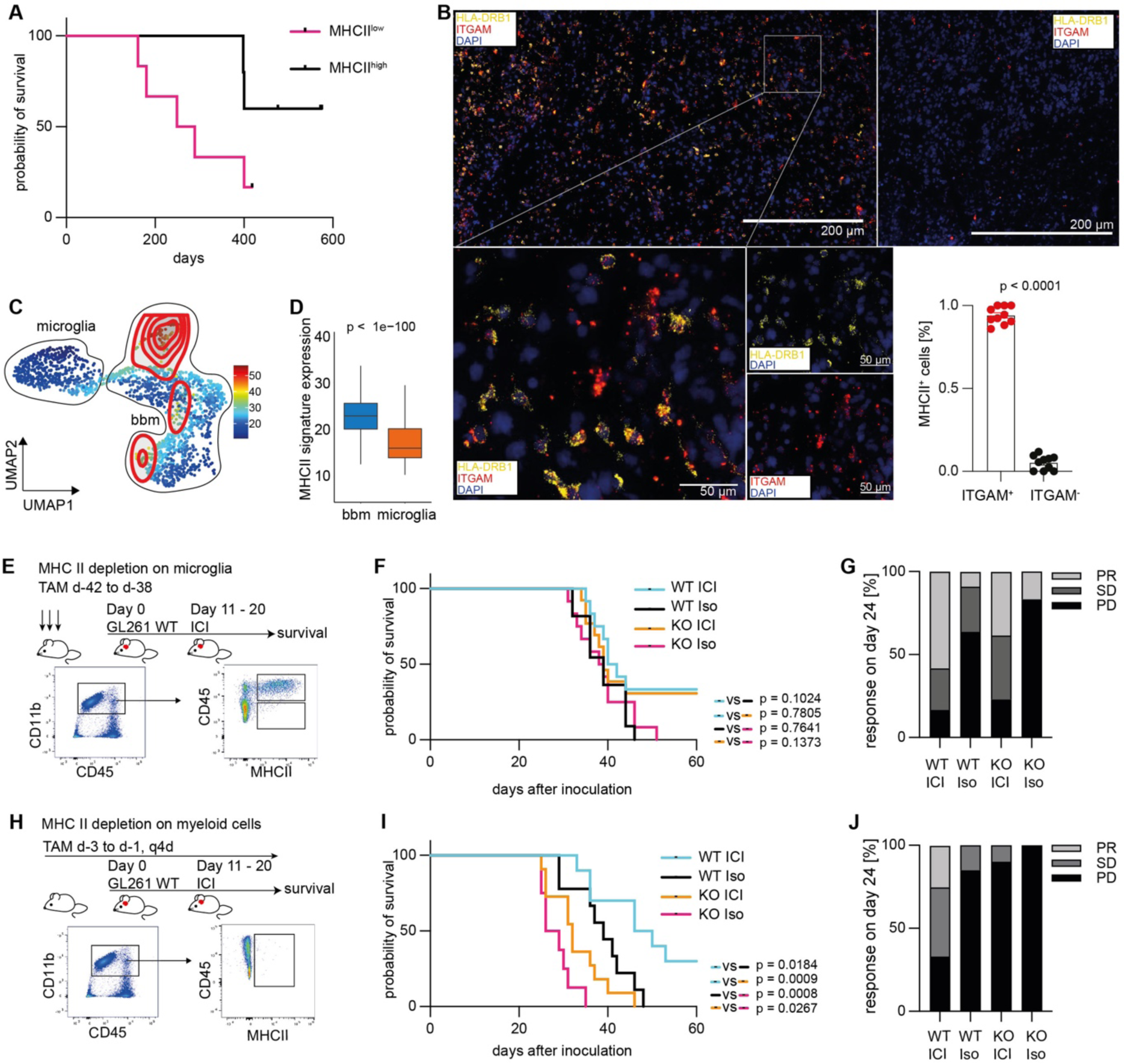
MHC class II expression on blood born tumor infiltrating myeloid cells is indispensable for immune checkpoint blockade and CD8^+^ T cell driven immune responses. (A) Overall survival of MHCII^high^ and MHCII^low^ IDH^wt^ patients treated with neoadjuvant αPD-1 blocking antibody (from (Cloughesy et al., 2019)). (B) In situ RNA hybridization in human GBM samples, n = 20 ROIs from n = 5 human GBM samples. (C) UMAP projection of antigen presentation signature. (D) Quantification of C. (E-F) Microglial MHCII knockout using timepoint-restricted dosing of tamoxifen 5 weeks before the experiment. Mice were treated with 5 doses αPD-1, αCTLA-4 and αPD-L1 every 72 h from day 12 after tumor inoculation. (E) Experimental overview. (F) KM survivor for MHCII^flox/flox^ (WT) and Cx3CR1CreERT2MHCII^flox/flox^ mice. (G) Distribution of partial responder (PR), stable disease (SD) and progressive disease (PD). (H-J) Myeloid MHCII knockout using continuous dosing of tamoxifen every 4-5 days. Mice were treated with 5 doses αPD-1, αCTLA-4 and αPD-L1 every 72 h from day 12 after tumor inoculation. (H) Experimental overview. (I) KM survivor for MHCII^flox/flox^ (WT) and Cx3CR1CreERT2MHCII^flox/flox^ mice. (J) Distribution of partial responder (PR), stable disease (SD) and progressive disease (PD). (B), (C) statistical significance was tested by unpaired t test. See also Figure S1, S2.

We then aimed at assessing the impact and mechanistic contribution of dynamic MHCII expression on microglia and bbm to brain tumor immunosurveillance. We generated a time-resolved tamoxifen-inducible MHCII knockout (MHCIIKO) mouse model by crossing Cx3cr1^CreERT2^ with MHCII^flox/flox^ mice, allowing for the discrimination of bbm, continuously replenished from the bone marrow, and long-living, tissue-resident microglia (Goldmann et al., 2013). MHCIIKO on microglia (microMHCIIKO) was achieved by time point-restricted administration of tamoxifen application 6 weeks before start of the tumor experiment, whereas MHCIIKO on myeloids (myeloidMHCIIKO) was induced by continuous tamoxifen administration (**Figure S2A-D**). MyeloidMHCIIKO Cx3cr1^CreERT2^-MHCII^flox/flox^ mice showed reduced CD4^+^ T cell abundance, but unperturbed frequencies in the myeloid, B cell and NK cell populations compared to control MHCII^flox/flox^ littermates (MHCIIWT) (**Figure S2E-H)**. We have previously shown that GL261 experimental gliomas respond to ICI in a T helper cell-dependent fashion (Aslan et al., 2020). In line with this and other reports (Reardon et al., 2016), MHCIIWT mice treated with αPD-1, αCTLA-4, and αPD-L1 showed radiographic responses and responding mice had improved overall survival (OS) when compared to vehicle treatment (**Figure 1E-J**). Surprisingly, microMHCIIKO did not affect response to ICI **(Figure 1F, G)**. In contrast, radiographic responses were abolished in mice displaying myeloidMHCIIKO (**Figure I, J**). We conclude that, in experimental gliomas, MHCII-restricted antigen presentation on bbm but not microglia is required for response to ICI.

### Myeloid MHCII expression drives cytotoxicity of tumor-reactive CD8^+^ T cells

Eradication of tumors is ultimately believed to be conferred by CD8^+^ T cell responses (Waldman et al., 2020). Consequently, in ICI-treated mice we observed a large increase of CD8^+^ T cell abundance compared to vehicle-treated mice (**Figure 2A**). Longitudinal scRNA-seq of CD45^+^ CD3^+^ murine tumor-infiltrating T cells revealed an increased expression of transcripts associated with CD8^+^ T cell effector phenotypes such as Pdcd1, Tnfrsf9, Gzmb and Prf1 at later timepoints compared to early-stage experimental gliomas (**Figure S3A-C**). Thus, in order to assess the impact of myeloidMHCIIKO on defined tumor-reactive CD8^+^ T cells we made use of the immunogenic OVA^257-264^ ^(SIINFEKL)^ MHCI-restricted antigen overexpressed in the GL261 experimental glioma model (GL261^SIINFEKL^). This resulted in an immunogenic cell line that was recognized and lysed by SIINFEKL-reactive OT-I T cells (**Figure 2B**, **Figure S3D**). GL261^SIINFEKL^-infiltrating CD8^+^ T cells were reactive against the SIINFEKL-peptide indicative of endogenous tumor-reactive CD8^+^ T cells in the TME (**Figure S1E**). As a result, GL261^SIINFEKL^ tumor growth was controlled in MHCIIWT mice. Similar to experimental observations in wildtype GL261-bearing mice, the radiographic control of GL261^SIINFEKL^ gliomas, however, was abolished in myeloidMHCIIKO mice and myeloidMHCIIKO mice had shortened overall survival (**Figure 2C, D, S3F**). Of note, tumor growth rapidly increased only after day 20, indicating a dynamic process of loss of tumor control. Flow cytometric phenotyping revealed no difference in intratumoral abundance of tumor-infiltrating CD8^+^ T cells in myeloidMHCIIKO animals, arguing against differential recruitment of CD8^+^ T cells into the tumor (**Figure 2E**). *Ex vivo* dextramer-staining of SIINFEKL-reactive CD8^+^ T cells enabled us to specifically interrogate tumor-reactive tumor-infiltrating cytotoxic T cells in myeloidMHCIIKO (**Figure 2F**). In myeloidMHCII-proficient and -deficient tumors, we did not observe differences in absolute numbers or fraction of SIINFEKL-reactive CD8^+^ T cells (**Figure 2F, S3G**). CD8^+^ T cells sorted from myeloidMHCIIKO GL261^SIINFEKL^ tumors, however, showed decreased killing capacity as compared to CD8^+^ T cells derived from myeloidMHCIIWT tumors (**Figure 2G**). In summary, intratumoral cytotoxic CD8^+^ T cell response but not CD8^+^ T cell recruitment to the tumor depends on MHCII-restricted antigen presentation by tumor-infiltrating bbm.

**Figure 2.**
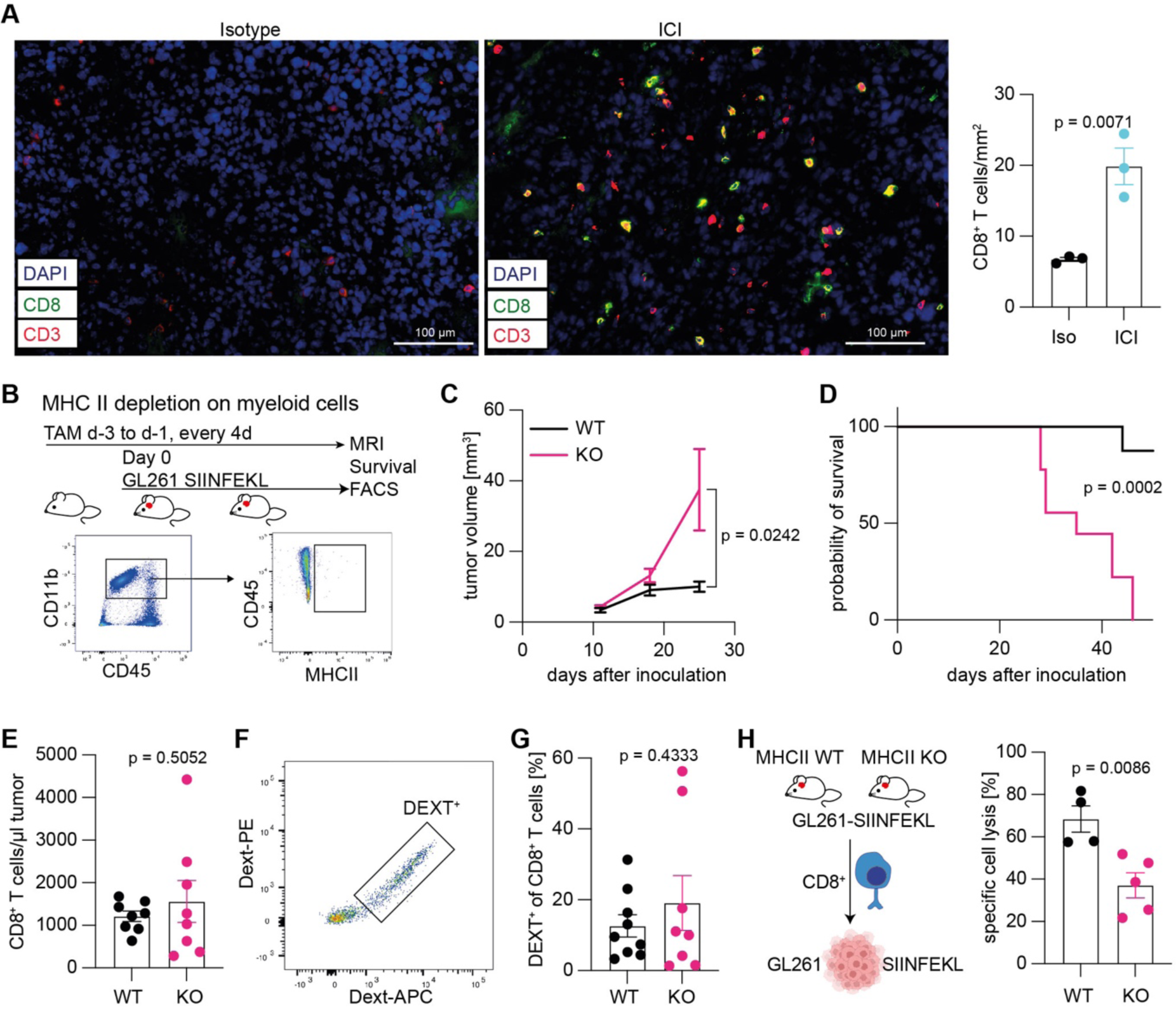
Myeloid MHCII expression drives cytotoxicity of tumor reactive CD8^+^ T cells. (A) Representative immunofluorescence image of isotype or ICI treated GL261 tumors in MHCIIWT mice stained for CD8 and CD3 (left). Quantification of CD8^+^ T cell infiltration (right). (B) Experimental overview for (C-G). I.c. injection of GL261^SIINFEKL^ tumor cells under continuous tamoxifen dosing. (C) Tumor volumes of GL261^SIINFEKL^ tumors in MHCII WT and MHCII KO mice measured by MR imaging. (D) Kaplan-Meier survival of tumor bearing mice. Statistical difference in survival was assessed by Mantel-Cox log-rank test. (E) CD8^+^ T cell infiltration of GL261^SIINFEKL^ tumors at day 20 after inoculation. (F) Representative flow cytometry gating for H-2K^b^-SIINFEKL dextramer stained CD8+ T cells (G) Percentage of H-2K^b^-SIINFEKL dextramer positive CD8+ T cells in GL261^SIINFEKL^ tumors at day 20 after inoculation. (H) Ex vivo GL261^SIINFEKL^ tumor cell lysis capacity of CD8^+^ T cells sorted from GL261^SIINFEKL^ tumors. Each symbol represents an individual mouse and statistical significance was tested by unpaired t test in (A), (C), (E), (G), (H). Bars indicate mean ± SEM. Dext, H-2K^b^-SIINFEKL dextramer; MFI, median fluorescence intensity. See also Figure S3.

### Loss of myeloid MHCII expression drives TOX-regulated dysfunctional CD8^+^ T cell states

The *ex vivo* persisting, reduced killing capacity in myeloidMHCII-deficient tumors is suggestive for a distinct cellular state of cytotoxic CD8^+^ T cells in the myeloidMHCIIKO TME. To identify underlying transcriptional programs in these CD8^+^ T cells, we performed scRNA-seq of CD8^+^ T cells isolated from GL261^SIINFEKL^ experimental gliomas at day 20 (**Figure 3A, S4A, B, C**). We identified 8 clusters that were transcriptionally defined by expression of stem-like and migratory genes like Tcf1, Klf2, expression of killer lectin-like genes like Klra1, Klrd1 and Klrb1c and genes associated with an activated phenotype like Gzma, Tigit, Pdcd1, Lag3 and Il10ra (**Figure S4C**). Differential gene expression (DGE) of CD8^+^ T cells revealed an upregulation of the interferon response gene Ifitm1 in myeloidMHCIIWT, whereas in myeloidMHCIIKO-derived CD8^+^ T cells Ccl5, Irf7 and the exhaustion markers Tox and Pdcd1 were highly upregulated (**Figure S4D**).

**Figure 3.**
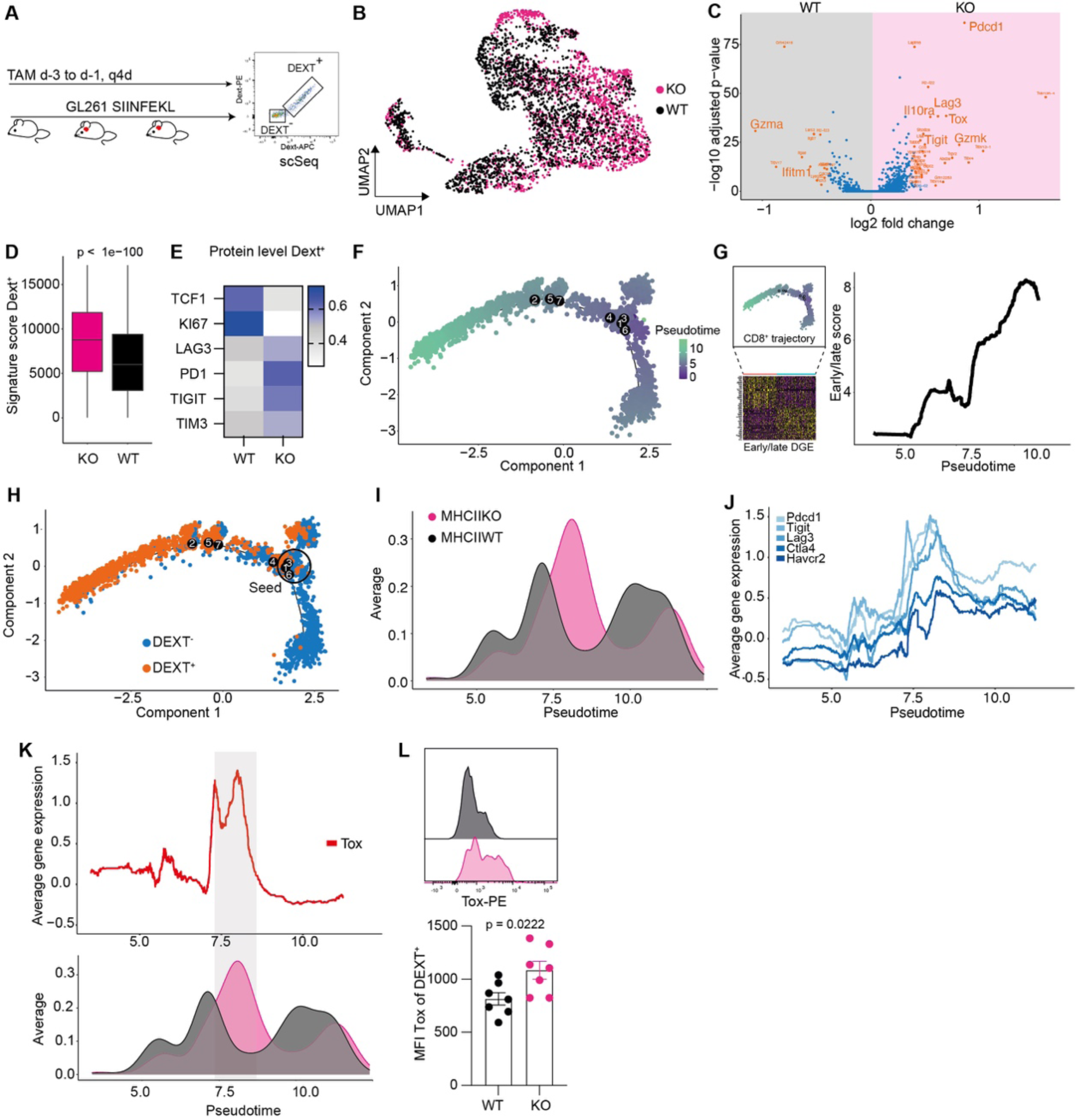
Loss of myeloid MHC class II expression drives tumor reactive T cells into distinct transcriptional exhaustion state. (A) Experimental overview. H-2K^b^-SIINFEKL dextramer positive (Dext^+^) and negative (Dext^-^) CD8^+^ T cells from GL261^SIINFEKL^ tumors from myeloidMHCIIKO and MHCIIWT littermates were sorted at day 20 after tumor inoculation and subsequently subjected to single-cell RNAseq. (B) UMAP visualization of single cell RNA-seq data, separated by genotype. (C) Volcano plot of differentially expressed genes in Dext^+^ CD8^+^ T cells from MHCIIWT and myeloidMHCIIKO mice. (D) Quantification of the dysfunctional signature in Dext^+^ CD8^+^ T cells MHCIIWT and myeloidMHCIIKO mice. Dysfunctional signature is composed of: Pdcd1, Lag3, Tigit, Ctla4, Havcr2, Tox. (E) Protein expression of exhaustion markers using flow cytometry. (F) Pseudotime trajectory analysis using monocle2 of CD8^+^ T cells. (G) Projection of early/late score from Figure S3 (A) on pseudotime of (F). (H) Trajectory analysis separated by Dext^+^ and Dext^-^ CD8^+^ T cells. (I) Average cell abundance of Dext^+^ CD8^+^ T cells from MHCIIWT and myeloidMHCIIKO mice along the pseudotime trajectory. (J) Rolling mean of exhaustion markers along the pseudotime trajectory in Dext^+^ CD8^+^ T cells. (K) Rolling mean of Tox expression along the pseudotime trajectory overlaid with average cell abundance. (L) TOX protein expression of Dext^+^ CD8^+^ T cells using flow cytometry, representative histogram (upper panel) and quantification (lower panel). (D) statistical significance was tested by unpaired t test. Each symbol represents an individual mouse and statistical significance was tested by unpaired t test in (L). Bars indicate mean ± SEM. Dext, H-2K^b^-SIINFEKL dextramer; MFI, median fluorescence intensity. See also Figure S4

To assess transcriptional T cell programs upon bbm MHCII deletion particularly in tumor-reactive CD8^+^ T cells, we performed scRNA-seq on dextramer-sorted SIINFEKL-reactive CD8^+^ T cells. DGE in tumor infiltrating Dext^+^ T cells indicated an upregulation of Gzma and Ifitm1 expression in MHCIIWT mice. In contrast, in myeloidMHCIIKO tumor-infiltrating Dext^+^ T cells displayed increased abundance of Pdcd1, Lag3, Tigit, Tox and Gzmk transcripts indicative of an exhausted phenotype (**Figure 3C**). Therefore, we defined a transcriptional and proteomic exhaustion signature including *Pdcd1, Tigit, Tim3, Lag3, Tox, Ctla4* (Li et al., 2019) enriched in myeloidMHCIIKO-retrieved SIINFEKL-reactive CD8^+^ T cells **(Figure 3D, E, S4E)**. Together with our observation that MHCIIKO-derived CD8^+^ T cells display reduced killing capacity (**Figure 2G**) we hypothesized that CD8^+^ T cells acquire a distinct dysfunctional transcription state in an MHCII-deficient TME. To assess gradual changes between MHCIIKO and MHCIIWT-derived CD8^+^ T cells we performed pseudotime analysis of SIINFEKL-reactive and bystander CD8^+^ (**Figure 3F**). Projection of an expression score derived from the longitudinal scRNA-seq of CD8^+^ T cells validated our pseudotime analysis and we observed reduced expression of stem-like associated genes like Tcf7, Ccr7 and Sell. (**Figure 3G, S4F**). Interestingly, SIINFEKL-reactive CD8^+^ primarily followed a different trajectory than bystander (non-SIINFEKL-reactive) CD8^+^ T cells indicative of their distinct state in the TME determined by their tumor-reactivity (**Figure 3H**). By calculating the normalized abundance of T cell states along the pseudotime trajectory of SIINFEKL-reactive CD8^+^ T cells retrieved from myeloidMHCII-proficient and -deficient microenvironments, we found an enrichment of a distinct T cell state in myeloidMHCIIKO CD8^+^ T cells that was defined by peak expression of exhaustion markers **(Figure 3I, J)**. Moreover, Tox expression peak robustly overlapped with myeloidMHCIIKO CD8^+^ enriched pseudotime-defined states (**Figure 3K**). Flow cytometric analysis of tumor-infiltrating Dext^+^ CD8^+^ T cells confirmed increased expression of TOX in MHCIIKO-derived cells (**Figure 3L**).

We therefore concluded that this exhausted TOX-regulated T cell state is dictated by the myeloid MHCII-deficient TME and that lack of MHCII expression on myeloids drives TOX-regulated dysfunctional CD8^+^ T cell states.

### Loss of MHCII drives chronic Nfat2 signaling and Tox-induced CD8 T cell dysfunction via osteopontin

CD4^+^ T cells license myeloid antigen-presenting cells in an MHCII-dependent manner (Ferris et al., 2020). However, in our model MHCII-proficient bbm displayed a bona fide immunosuppressive signature defined by differential upregulation of *Cd274, Mrc1* and *Arg1* transcripts and a reduced co-stimulatory signature defined by CD86, CD80, Icoslg and Tnfsf4 compared to MHCII-deficient bbm arguing against an MHCII-CD4 dependent licensing of antigen presenting cells (**Figure 4A**). Of note, upregulation of negative regulators of immune function such as immune checkpoint molecules are signified by feedback loops resulting from preceding pro-inflammatory stimuli. Consequently, in line with previous reports, bypassing CD4^+^ T cell-mediated activation of the myeloid compartment via MHCII using a CD40-agonistic antibody did not lead to improved but rather diminished tumor control in myeloidMHCIIKO mice (**Figure S5A**) (van Hooren et al., 2021).

**Figure 4.**
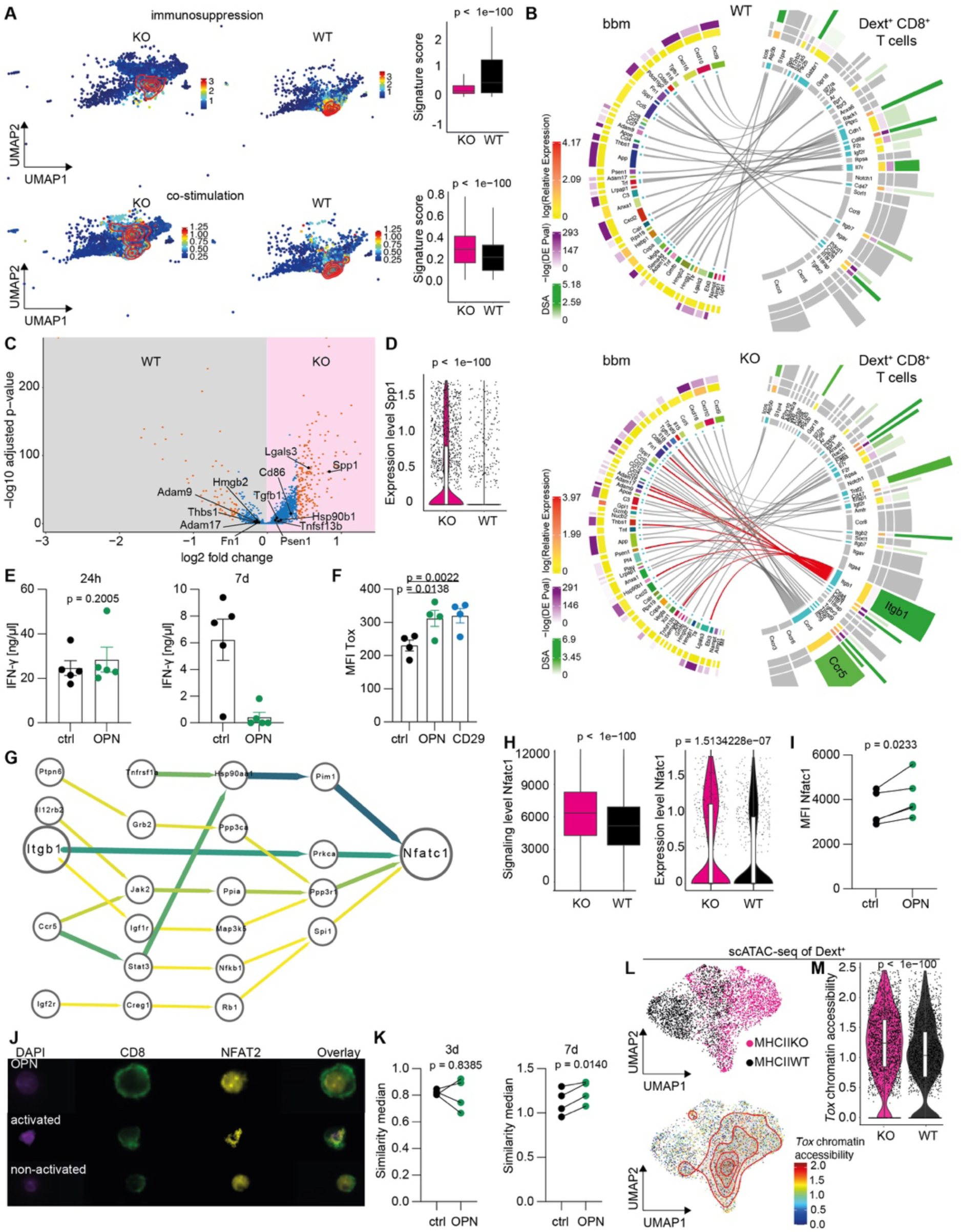
OPN on bbm leads to chronic Nfat2 translocation and Tox-mediated CD8^+^ T cell dysfunction (A) Density plot of immunosuppressive and co-stimulatory signature on the myeloid compartment of GL261^SIINFEKL^-derived CD45^+^ cells separated by genotype (left panel) and quantification (right panel). (B) Ligand receptor interaction map between bbm and Dext^+^ CD8^+^ T cells from MHCIIWT and MHCIIKO mice. (C) Volcano plot of differentially expressed genes in bbm from MHCIIWT and MHCIIKO mice. (D) Expression of SPP1 in bbm. (E) IFN-γ release of OT-I splenocytes stimulated with SIINFEKL-peptide measured by ELISA 24h and 7d after first activation for 24h each. Values below detection limit were set to the minimum values for visualization. (F) MFI of Tox in SIINFEKL activated OT-I T cells (upper panel) and number of CD8^+^ T cells in wells treated with OPN or activating CD29 antibody 7 days after activation (G) Upstream signaling of Nfatc1 in Dext^+^ CD8^+^ T cells from MHCIIKO mice. (H) Signaling level of Nfatc1 (left) and expression of Nfatc1 (right) in Dext^+^ CD8^+^ T cells. (I) MFI of NFAT2 in OT-I T cells treated with OPN 7 days after activation. (J) Representative Imagestream analysis of OT-I T cells 7 days after activation (purple DAPI, green CD8, yellow NFAT2) (K) Quantification of NFAT2 translocation 3 and 7 days after priming. (L) UMAP projection of single cell ATAC sequencing of Dext^+^ CD8^+^ T cells (upper panel), density plot of Tox accessibility (lower panel). Bars indicate mean ± SEM. Statistical significance was tested by unpaired t test. (M) Quantification of (L). See also Figure S5. Statistical significance was tested by unpaired t test in (A), (D), (H), (M). (E), (F) bars indicate mean ± SEM. Statistical significance was tested by paired t test in (E), (I), (K). For (E) 7d no statistical testing was applied as values were below the detection limit. Statistical significance was tested by RM one-way ANOVA with Tukey’s test in (F). See also Figure S5.

To characterize the cell - cell interaction between myeloidMHCII with CD8^+^ T cells we again performed receptor ligand analyses between bbm and SIINFEKL-reactive CD8^+^ T cells from GL261^SIINFEKL^ tumors. We found increased predicted interactions via *Itgb1* and *Ccr5* in the myeloidMHCII-deficient but no increased interactions in the myeloidMHCII proficient microenvironment (**Figure 4B**). *Itgb1* encodes for the cell surface receptor CD29. CD29 has been previously reported as surrogate marker for highly cytotoxic and IFN-ɣ-producing human T cells (Nicolet et al., 2020) and is upregulated in activated murine T cells (**Figure S5B)**. One of its predicted ligands, osteopontin (OPN) which is transcribed by SPP1, was highly upregulated in MHCIIKO bbm (**Figure 4C, D**). To investigate if OPN can induce CD8^+^ T dysfunction we treated OT-I cells with OPN. Peptide stimulation of OT-I T cells in presence of OPN did not lead to differences in IFN-γ release in the initial response. However, after 7 days of peptide exposure, we observed strong decrease of IFN-γ release in OPN treated OT-I T cells. (**Figure 4E**). In our murine myeloidMHCIIKO model, CD8^+^ T cell dysfunction was associated with To*x* expression. Indeed, *in vitro* OPN increased TOX protein expression in activated OT-I T cells as well as an agonistic αCD29 antibody (**Figure 4F**).

Next, we aimed to investigate how OPN induces Tox expression in activated CD8^+^ T cells. Like *Ptprc* encoding for CD45, the OPN ligand CD29 has been implied in increased Akt signaling resulting in enhanced T cell receptor signaling followed by NFAT transcriptional activity (Jiang et al., 2015; Saunders and Johnson, 2010). NFAT activation has recently been shown to mediate Tox expression and T cell exhaustion (Khan et al., 2019; Scott et al., 2019). Strikingly, upstream pathway analyses of Nfatc1, encoding for NFAT2 in SIINFEKL-reactive CD8^+^ T cells in the context of myeloidMHCII deficiency predicted Itgb1 in Nfatc1 upstream signaling (**Figure 4G**). In our pseudotime analysis, Nfatc1 expression coincided with Tox expression and Nfatc1 and its downstream signaling pathway were upregulated in SIINFEKL-reactive CD8^+^ T cells derived from myeloidMHCIIKO tumors (**Figure 4H, S5C**). *In vitro* OPN treatment increased Nfatc1 expression following T cell activation (**Figure 5I**). Transcription factors like NFAT2 are not only transcriptionally regulated but are in steady-states inactively localized in the cytoplasm and translocate into the nucleus upon activation. Therefore, we also assessed NFAT2 translocation using imaging flow cytometry. Indeed, we observed significantly increased NFAT translocation in OT-I T cells 7 days after activation (**Figure 4J, K**). More recently, chronic Nfat activation been reported to increase accessibility of Tox (Khan et al., 2019; Scott et al., 2019). Conclusively, scATAC-seq of SIINFEKL-reactive CD8^+^ T cells revealed an increased chromatin accessibility of *Tox* in myeloidMHCIIKO-retrieved compared to control SIINFEKL-reactive CD8^+^ T cells (**Figure 4L, M, S5D**).

**Figure 5.**
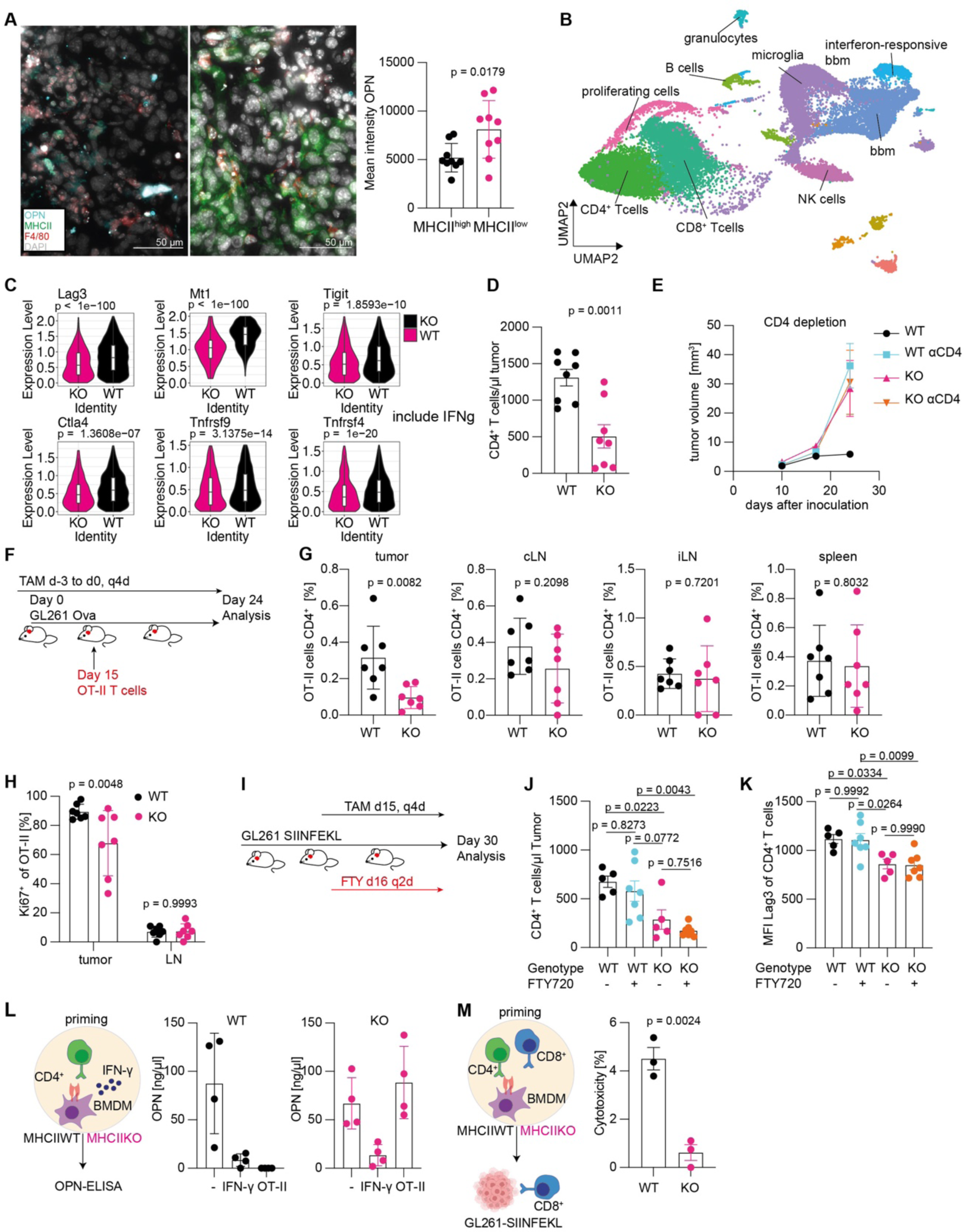
Intratumoral MHCII presentation and CD4^+^ T cell limits OPN expression on bbm. (A) Representative immunofluorescence image of F4/80 (red), MHCII (green) and OPN (cyan) in GL261^SIINFEKL^ tumors (left) and quantification (right). See also Figure S6A. (B) Visualization of scRNA-seq data using UMAP of CD45^+^ GL261^SIINFEKL^ tumors at day 20. (C) Expression of activation markers on CD4^+^ T cells from (B). (D) Abundance of CD4^+^ T cells in GL261^SIINFEKL^ tumors derived from MHCII KO and WT mice, measured by flow cytometry. (E) Growth curves of GL261^SIINFEKL^ tumors after antibody mediated CD4^+^ T cell depletion in MHCIIWT and myeloidMHCIIKO mice. (F) Experimental overview. OT-II T cells were adoptively transferred in GL261^Ova^ bearing myeloidMHCIIKO and MHCIIWT mice and subsequently analyzed using flow cytometry. (G) Abundance of transferred OT-II T cells in tumor and secondary lymphoid organs analyzed using flow cytometry. cLN cervical lymph nodes, iLN inguinal lymph node (H) Ki67 expression of transferred OT-II T cells in tumor and cervical lymph nodes. (I) Experimental overview. GL261^SIINFEKL^ tumors were injected and myeloidMHCIIKO was induced in mice with established tumors at day 15, with subsequent continuous FTY720 treatment. Tumor infiltrating immune cells were analyzed on day 30. (J) Absolute numbers of tumor infiltrating CD4^+^ T cells. (K) MFI of LAG3 of tumor infiltrating CD4^+^ T cells. (L) OPN concentration in the supernatant of BMDMs from MHCIIWT or MHCIIKO mice treated with 20 ng/µl IFN-γ or OT-II T cells and OVA^323-339^ peptide. Values below the detection limit were set to the minimum detection limit for visualization. No statistic testing was applied. (M) Cytotoxicity of OT-I T cells after priming with SIINFEKL-peptide loaded BMDMs from MHCIIWT and MHCIIKO mice co-cultured with Ova^323-339^ activated OT-II T cells. For (A), (D), (G), (H), (J), (K) bars indicate mean ± SEM. Statistical significance was tested by unpaired t test in (A), (C), (D), (G), (H), (M). Statistical significance was tested by one-way ANOVA with Tukey’s test in (J), (K). See also Figure S6.

Taken together, our results indicate that OPN released by MHCII-deficient bbm activates CD29 on CD8^+^ T cells, resulting in prolonged Nfat2 signaling and eventually, Tox-mediated CD8^+^ T cell dysfunction.

### Intratumoral MHCII-restricted antigen presentation and CD4^+^ T cell activation is required for functional cytotoxic T cell states

We next sought to investigate how MHCII regulates OPN expression in the tumor microenvironment. In GL261^SIINFEKL^ tumors OPN was primarily found in MHCII^low^ regions suggesting a direct relationship between OPN expression and MHCII abundance (**Figure 5A, S6A**). To interrogate myeloidMHCIIKO-associated microenvironmental alterations that drive cytotoxic T cell exhaustion in GL261^SIINFEKL^ tumors, we integrated scRNA-seq from sorted CD45^+^ CD3^-^ myeloid cells and CD3^+^ Dex^SIINFEKL^-negative T cells including CD4^+^ T cells from myeloidMHCII-proficient and -deficient GL261^SIINFEKL^ tumors (**Figure 5B, S4A)**. ScRNA-seq-based interaction analysis revealed reduced interactions between CD4^+^ T cells and bbm in the MHCII-deficient TME. Specifically, we found CD4^+^ - bbm interactions for Csf2ra, Ifnar2, Tlr4 and several integrins (Itga5, Itgb5, and Itgb1) exclusively predicted in MHCIIWT tumors, suggesting intratumoral cell - cell interactions between CD4^+^ T cells and MHCII-proficient bbm that define bbm phenotype (**Figure S6B**). In myeloidMHCII-deficient tumors, we observed reduced CD4^+^ T cell numbers and CD4^+^ T cells displayed reduced expression of activation markers including reduced expression of Ifng (**Figure 5** **C, D)**. Depletion of CD4^+^ T cells mimicked the myeloidMHCIIKO setting as control of GL261^SIINFEKL^ tumors was abolished in CD4^+^ T cell-depleted animals (**Figure 5E**). Of note, the tamoxifen-inducible myeloidMHCIIKO mouse model does not allow to discriminate between intratumoral and extracerebral MHCII-restricted antigen presentation. To decipher the relevant compartment for MHCII-restricted antigen presentation, we additionally made use of the GL261^OVA^ tumor model to identify T cell responses to a tumor-derived MHCII-restricted antigen. Using adoptive transfer of OT-II T cells specifically recognizing the immunodominant MHCII-restricted OVA^323-339^ epitope, we found antigen-restricted effector cytokine production and CD4^+^ T cell proliferation in dependence of myeloidMHCII-proficiency exclusively in the experimental GL261^OVA^ tumor but not in cervical or inguinal lymph nodes nor the spleen (**Figure 4F-H**). Additionally, we only observed endogenous OVA^323-339^-reactive CD4^+^ T cells in the TME (**Figure S6C**). Next, we inhibited T cell infiltration from lymphoid organs into established tumors using fingolimod (FTY720) in myeloidMHCIIKO mice (**Figure 4I**). As expected, FTY720 treatment led to decreased T cell numbers in the TME (**Figure S6D**). After 14 days of treatment, we observed decreased abundance of CD4^+^ T cells and decreased expression of the activation marker Lag3 in FTY720-treated myeloidMHCIIKO mice compared to MHCIIWT littermates, arguing for an intratumoral MHCII-dependent proliferation and activation of CD4^+^ T cells (**Figure 4J, K, S6E**). Next, in order to validate the functional relationship between MHCII and OPN, we treated *in vitro*-generated MHCIIWT and MHCIIKO bone marrow derived macrophages (BMDM) with IFN-γ or co-cultured the latter with OVA^323-339^-activated OT-II T cells. IFN-γ-treatment and activated OT-II cells in particular reduced OPN expression below the detection limit in an MHCII-proficient co-culture (**Figure 5L**). Thus, we concluded that IFN-γ released by intratumorally MHCII-restricted antigen-activated CD4^+^ T cells reduces OPN expression in tumor infiltrating bbm. Consequently, peptide-activated CD8^+^ OT-I T cells co-cultured with BMDMs and activated OT-II cells displayed increased cytotoxicity and decreased TOX expression in an MHCII-proficient and therefore OPN-reduced condition (**Figure 5M, S6F, G)**.

This observation suggests that in our model intratumoral MHCII-restricted antigen presentation to CD4^+^ T cells suppresses OPN expression and maintains a functional cytotoxic CD8^+^ T cell pool.

### MHCII on blood-borne myeloids is associated with exhaustive CD8^+^ T cell states in gliomas

To study functional implications of myeloid MHCII expression on cytotoxic CD8^+^ T cell states in human gliomas, we performed scRNA-seq of CD45^+^ CD3^+^ and CD45^+^ CD3^-^ cells from 9 patients with malignant glioma (**Figure 6A, C, S7A, B, E**). Similar to observations in our murine experimental systems, we found two exhausted CD8^+^ T cell states signified by *TOX, PDCD1, TIGIT, LAG3, CTLA4, and HAVCR2* expression and high *ITGB1* expression (**Figure 6B, S7C, D)**. For TMEM119^+^ microglia as well as bbm, we identified two clusters with differential MHCII-associated gene signatures (**Figure 6C, S7F)**. During glioma progression, the fraction of bbm within the TME increases (Friebel et al., 2020; Friedrich et al., 2021; Klemm et al., 2020). Hence, the proportion of bbm within the TME is dependent of disease duration, treatment and date of diagnosis. To comparatively assess MHCII expression on bbm in relation to CD8^+^ T cells we calculated the proportion of bbm^MHCIIlow^ to bbm^MHCIIhigh^ per patient. Strikingly, we found a negative correlation between MHCII expression on bbm and the intratumoral dysfunctional state of CD8^+^ T cells **(Figure 6D, E)**. Of note, SPP1 expression negatively correlated with MHCII expression our single cell dataset in MHCII^high^ bbm and across patients as well as with MHCII expression normalized to CD14 expression in the bulk RNAseq dataset derived from the TCGA GBM cohort (**Figure 6F, G, S7G**). Stem-like progenitor exhausted CD8^+^ T cells expressing TCF1 have been shown to drive response to ICI in human and murine solid tumors (Kurtulus et al., 2019; Miller et al., 2019; Sade-Feldman et al., 2018; Siddiqui et al., 2019). In human GBM tissue samples, we observed accumulation of TCF1^+^ CD8^+^ T cells in the proximity of MHCII^+^ ITGAM^+^ myeloid cells. In contrast, in MHCII^low^ niches, we observed enriched HAVCR2^+^ CD8^+^ T cells, indicative of an exhaustive phenotype (**Figure 6H**).

**Figure 6.**
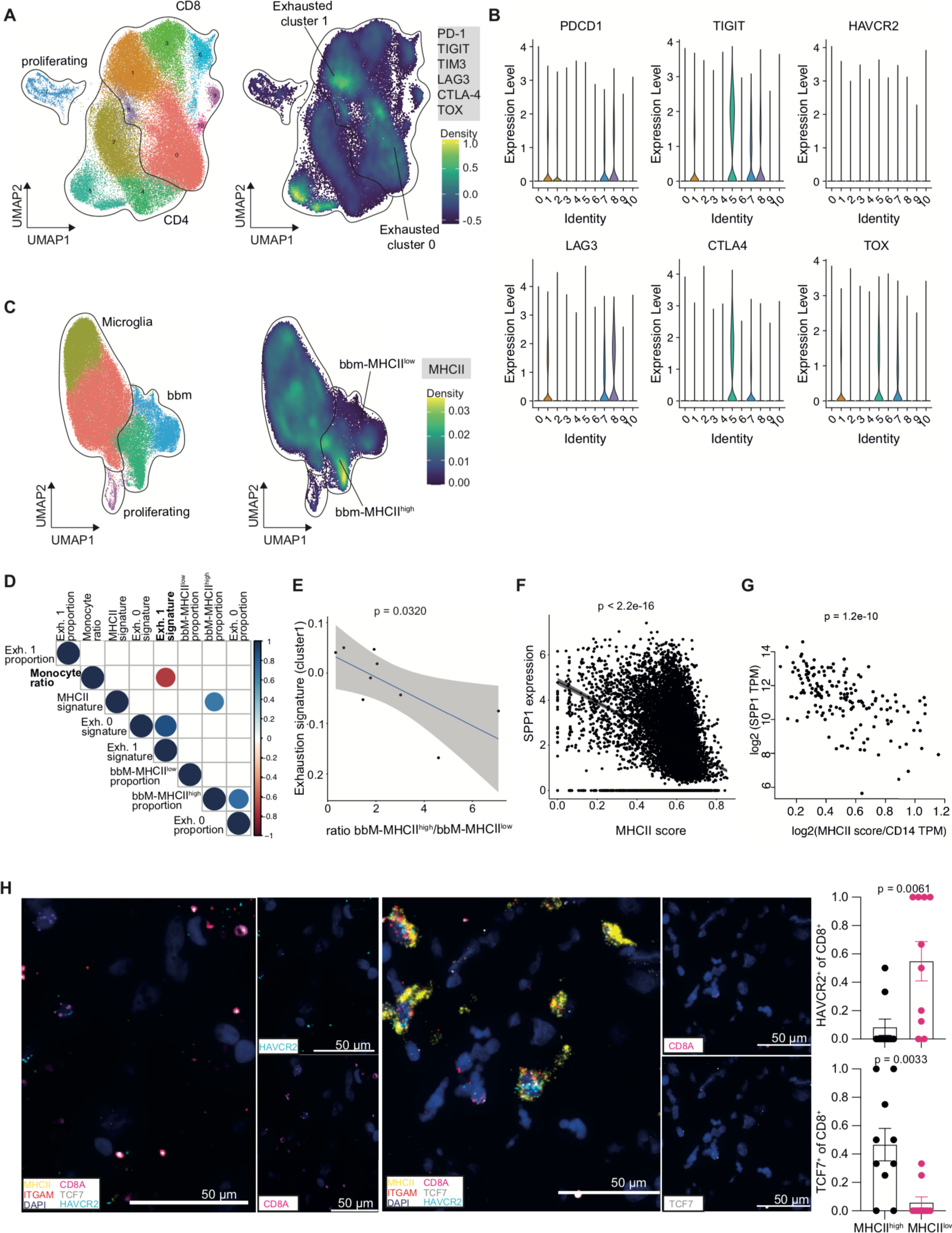
Lack of MHC class II drives CD8^+^ T cell exhaustion in human glioblastoma. (A-F) Single cell RNAseq of human GBM infiltrating immune cells. n = 9. (A) UMAP of T cells, density plot of dysfunction signature. (B) Violin plots of exhaustion markers in clusters defined in (A). (C) UMAP of myeloid cells, density plot of MHCII-signature from Figure 1C. (D) Correlation plot of immune cell subsets and exhaustion signatures, significant correlations are shown. (E) Correlation of the ratio of MHCII^high^/MHCII^low^ bbms and exhaustive T cell state in cluster 1. (F) Correlation of MHCII expression and SPP1 expression in GBM derived bbms. (G) Pearson correlation of SPP1 expression and MHCII signature expression normalized to CD14 expression in the TCGA GBM dataset. (H) In situ hybridization in human GBM tissue using probes against MHCII, ITGAM, CD8A, TCF7 and HAVCR2. Representative images for MHCII^low^ (left) and MHCII^high^ (middle) regions are shown. Right: Quantification. n = 20 ROIs of 5 GBM samples. Statistical significance was tested by unpaired t test. See also Figure S7.

In summary we demonstrate by combined T cell and myeloid scRNA-seq that in human gliomas, reduced MHCII expression on bbm is correlated with increased OPN expression and exhaustive intratumoral CD8^+^ T cell states corroborating our mechanistic studies in experimental mouse systems.

## Discussion

The infiltration of tolerogenic and immunosuppressive myeloid cells is a hallmark of disease progression in brain tumors (Friebel et al., 2020; Friedrich et al., 2021; Hambardzumyan et al., 2015; Klemm et al., 2020; Pathania et al., 2017). While previous therapeutic strategies have focused on the depletion of these cell types, there is now increasing efforts in reprogramming immunosuppressive microenvironments (Coniglio et al., 2012; Pyonteck et al., 2013; Quail and Joyce, 2017). These reprogramming strategies have largely focused on skewing the phenotype of bbm from an immunosuppressive M2-like to a proinflammatory M1-like phenotype assuming that immunosuppression by bbm is driven by soluble factors such as IL10 or TGFb (Poon et al., 2017; Ravi et al., 2022). MHCII expression is considered a marker of proinflammatory M1-like bbm but the functional relevance of MHCII has remained ill-understood. Here we provide evidence that blood-borne myeloids (bbm) represent a critical source of intratumoral MHCII-restricted antigen presentation to maintain stemness of cytotoxic T cells in brain tumors.

We provide an additional mechanistic layer by dissecting complex cell-cell interaction loops that can prevent CD8^+^ T cell exhaustion. The presence of macrophages in the TME often correlates with poor prognosis in various cancer entities (Noy and Pollard, 2014; De Palma and Lewis, 2013). Notably, in our setting, MHCII-proficiency on bbm was indispensable for anti-tumor activity, even though MHCII-proficient bbm displayed a *bona fide* immunosuppressive and a reduced co-stimulatory phenotype. Inflammatory stimuli often lead to feedback loops triggering expression of immunosuppressive and checkpoint molecules. This immunosuppressive phenotype in our setting in established tumors seems to be uncoupled from a functional cytotoxic CD8^+^ T cell state and ICI-induced anti-tumor response. Additionally, in line with other reports, our results suggest that a CD8^+^ T cell response to ICI requires a functional activated state of intratumoral T cells that can be reinvigorated (Im et al., 2016; Miller et al., 2019).

MHCII-restricted neoantigens have previously been shown to be a critical requirement for anti-tumor responses following immunotherapy (Alspach et al., 2019; Kreiter et al., 2015). CD4^+^ T cell responses induced by MHCII-epitope complexes form a prerequisite for maintaining cytotoxic CD8^+^ T cell responses in tumors and viral infections (Ahrends et al., 2017, 2019; Zander et al., 2019). Based on these observations, CD4^+^ T cells have been shown to license antigen presenting cells in a CD40-dependent manner in tumor draining lymph nodes that subsequently prime CD8^+^ T cells for efficient anti-tumor activity (Binnewies et al., 2019; Ferris et al., 2020). However, in our study, and in line with previous reports, bypassing APC licensing using agonistic CD40 antibody did not result in improved priming of MHCII-derived tumor-reactive CD8^+^ T cells arguing for a CNS-specific mechanism (van Hooren et al., 2021). Furthermore, we showed that the loss of intratumoral MHCII-restricted antigen presentation results in loss of tumor control by CD8^+^ T cells in brain tumors. This loss of tumor control was associated with a dysfunctional CD8^+^ T cell state orchestrated by increased Tox expression. In human GBM samples, we found progenitor-exhausted functional CD8^+^ T cells in the proximity of intratumoral myeloid-MHCII^+^ subniches, indicative of a MHCII-dependent maintenance of a functional cytotoxic T cell pool. Similar observations have been made in kidney cancer, and T cell attracting myeloid immune cell hubs have recently been described in colorectal tumors, indicating that spatially organized immune cell clusters are required for intratumoral immune cell crosstalk, and ultimately, the maintenance of functional anti-tumor responses (Jansen et al., 2019; Pelka et al., 2021). Others have recently described a spatial co-dependency of intratumoral macrophages and exhausted CD8^+^ T cells in hypoxic tumor regions (Kersten et al., 2021).

Mechanistically, we have shown that OPN, locally released by intratumoral myeloid cells, induces chronic NFAT2 translocation in tumor reactive CD8^+^ T cells. Chronic NFAT2 activation, as recently described (Khan et al., 2019; Scott et al., 2019), eventually leads to TOX-mediated dysfunctional transcriptional state of CD8^+^ T cells that was associated with loss of tumor control. In line with this, Klement et al. have recently shown that high concentration of OPN reduces T cell activation and proliferation (Klement et al., 2018). OPN has been described in different tumor entities to constitute a negative prognostic factor that also correlates with glioma grade (Sreekanthreddy et al., 2010; Toy et al., 2009; Weber et al., 2010). Mechanistically, excessive release of OPN recruits M2 macrophages to the TME that impair anti-tumor immunity (Wei et al., 2019). By contrast, in our study, we found that bbm provide a relevant source of OPN that directly impairs cytotoxic CD8^+^ T cell functions by driving chronic activation after antigen-recognition. We show that the intratumoral activation of CD4^+^ T cells drastically reduces local OPN accumulation via INF-γ release. Thus, therapeutic modulation of the TME by targeting MHCII-restricted neoepitopes via CD4^+^-specific immunotherapy might enable durable tumor control by cytotoxic CD8^+^ T cells (Kilian et al., 2021a; Poncette et al., 2022). Adoptive T cell therapies are currently being investigated for the treatment of GBM (Kilian et al., 2021b; Larson et al., 2022; Majzner et al., 2022). The necessity of combined CD4 and CD8-lineage determined T cellular products for an efficacious therapeutic regime has recently been demonstrated (Sommermeyer et al., 2016). Defining the 1:1 ratio of CD4^+^:CD8^+^ CAR T cells confers superior antitumor reactivity in vivo, indicating the synergistic antitumor effects of the two subsets (Turtle et al., 2016). Consequently, pharmacological suppression of the OPN-CD29-NFAT-TOX axis might mimic synergistic CD4^+^ activation in eliciting full-fledged antitumor responses, when MHC-II antigen-presentation or functional CD4^+^ T cells are limited.

In summary, our study reveals a critical role of intratumoral MHCII-dependent antigen presentation in brain tumors in modulating the TME and preventing terminal dysfunctional states of cytotoxic T cells. These findings provide a basis for the rational design of CD8^+^ T cell-targeting immunotherapies against brain tumors. Our data further suggest that response to ICI in the CNS is limited to myeloidMHCII-proficient microenvironments and that modulating OPN expression might synergize with immunotherapy.

## Author contributions

M.K. conceptualized the study, designed and performed experiments, analyzed and interpreted data, and wrote the paper. R.S. analyzed scRNA-seq and scATAC data, interpreted data, and wrote the paper. C.L.T. analyzed scRNA-seq data. C.K., M.F., K.S. and F.C. performed in vitro and in vivo experiments. A.K. participated in data interpretation. K.L. processed human tissue and generated human single cell RNA libraries. S.J. and K.J. performed in vitro experiments and generated murine single cell RNA and ATAC libraries. M.R. provided GBM tissue and initial analysis of GBM samples. R.P., A.vD., and W.W. were involved in data interpretation. A.M., L.B. and M.P. conceptualized and supervised the study, interpreted data and wrote the paper.

## Acknowledgements

Acknowledgements and funding

We acknowledge the support of the DKFZ Light Microscopy Facility, the DKFZ Genomics and Proteomics Core Facility, the Center for Preclinical Research, the Flow Cytometry Core Facility and the single-cell Open Lab at the German Cancer Research Center. We acknowledge the data storage service SDS@hd supported by the Ministry of Science Germany. This study was supported by grants from the Else Kröner Fresenius Foundation, the Swiss Cancer Foundation, the University Heidelberg Foundation, the DFG (German Research Foundation) – Project-ID 404521405, SFB 1389 - UNITE Glioblastoma B03 to L.B., the Deutsche Forschungsgemeinschaft (DFG, German Research Foundation, projects 404521405 (CRC1389 “Understanding and targeting resistance in glioblastoma, Work Packages B01 and B02) to M.P. A.M. was supported by The Alon fellowship for outstanding young scientists, Israel Council for Higher Education, from the Israel Science Foundation (1700/21), from the Israel Cancer Association (ICA, 01028753) and from the Israel Cancer Research Fund (ICRF) Research Career Development Awards. M.K. received fellowships by the Helmholtz International Graduate School for Cancer Research and the Landesgraduiertenförderung (LGF). R.S was supported in part by a fellowship from the Edmond J. Safra Center for Bioinformatics at Tel-Aviv University.

## KEY RESOURCES TABLE

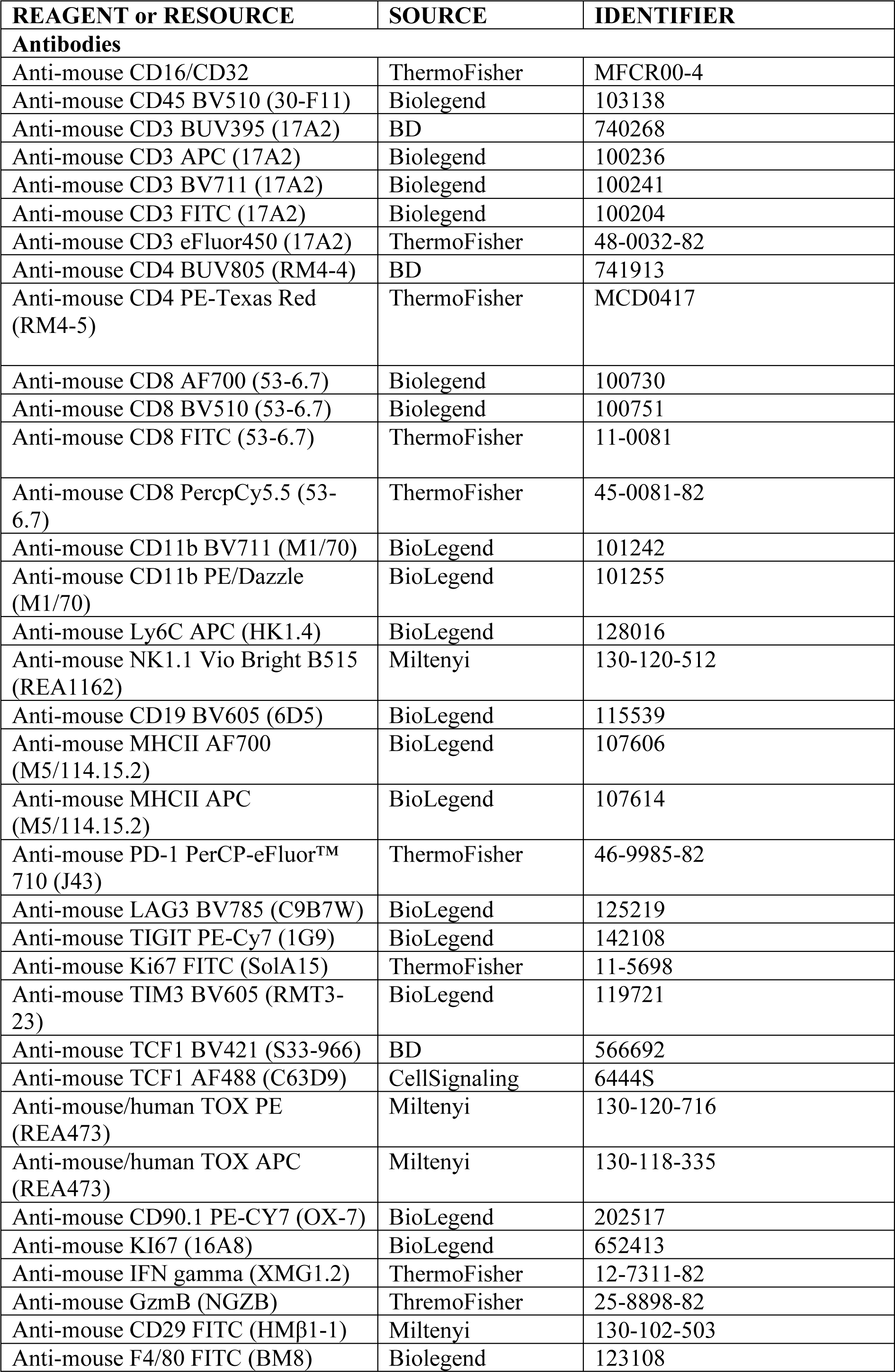
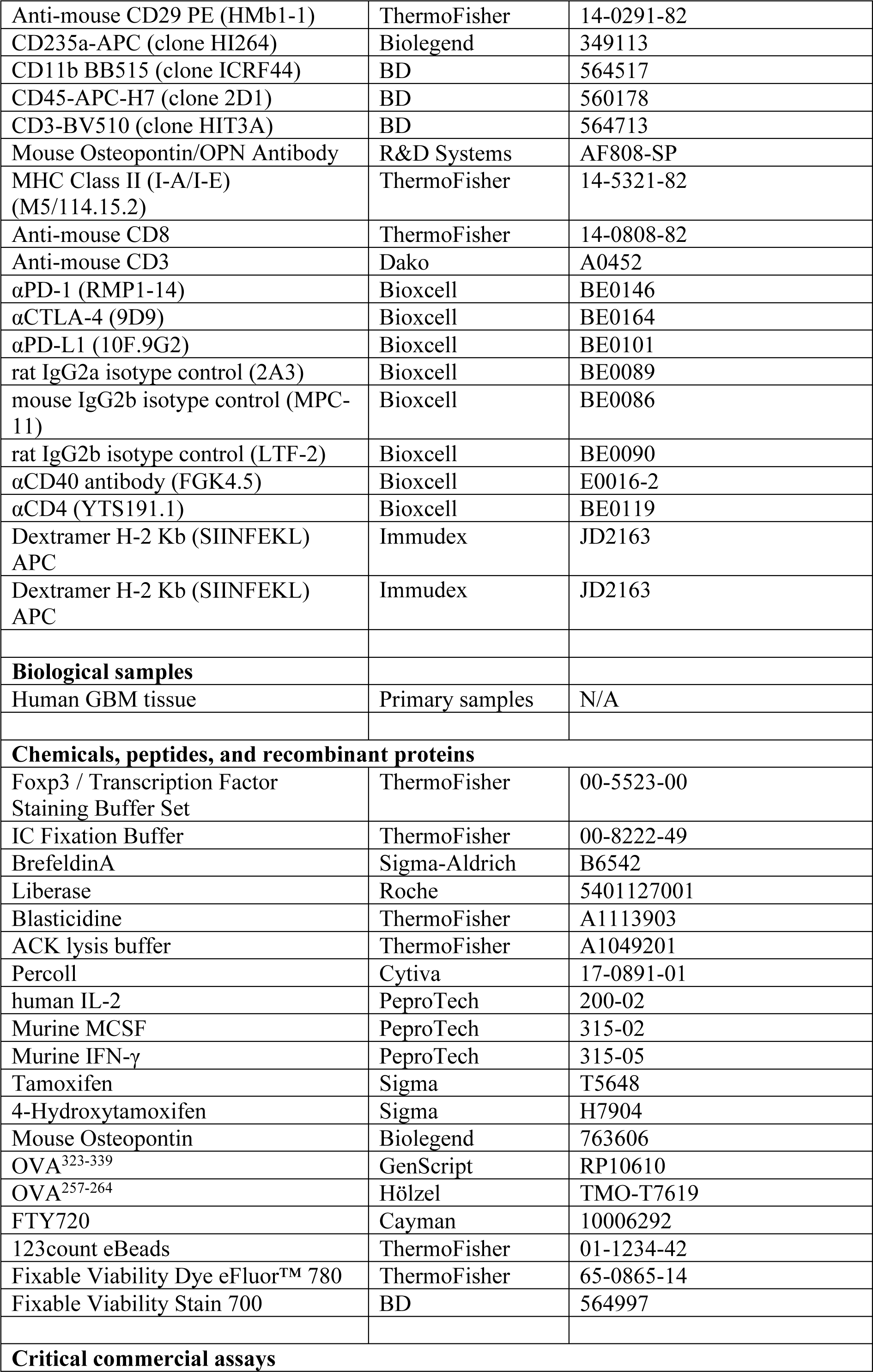
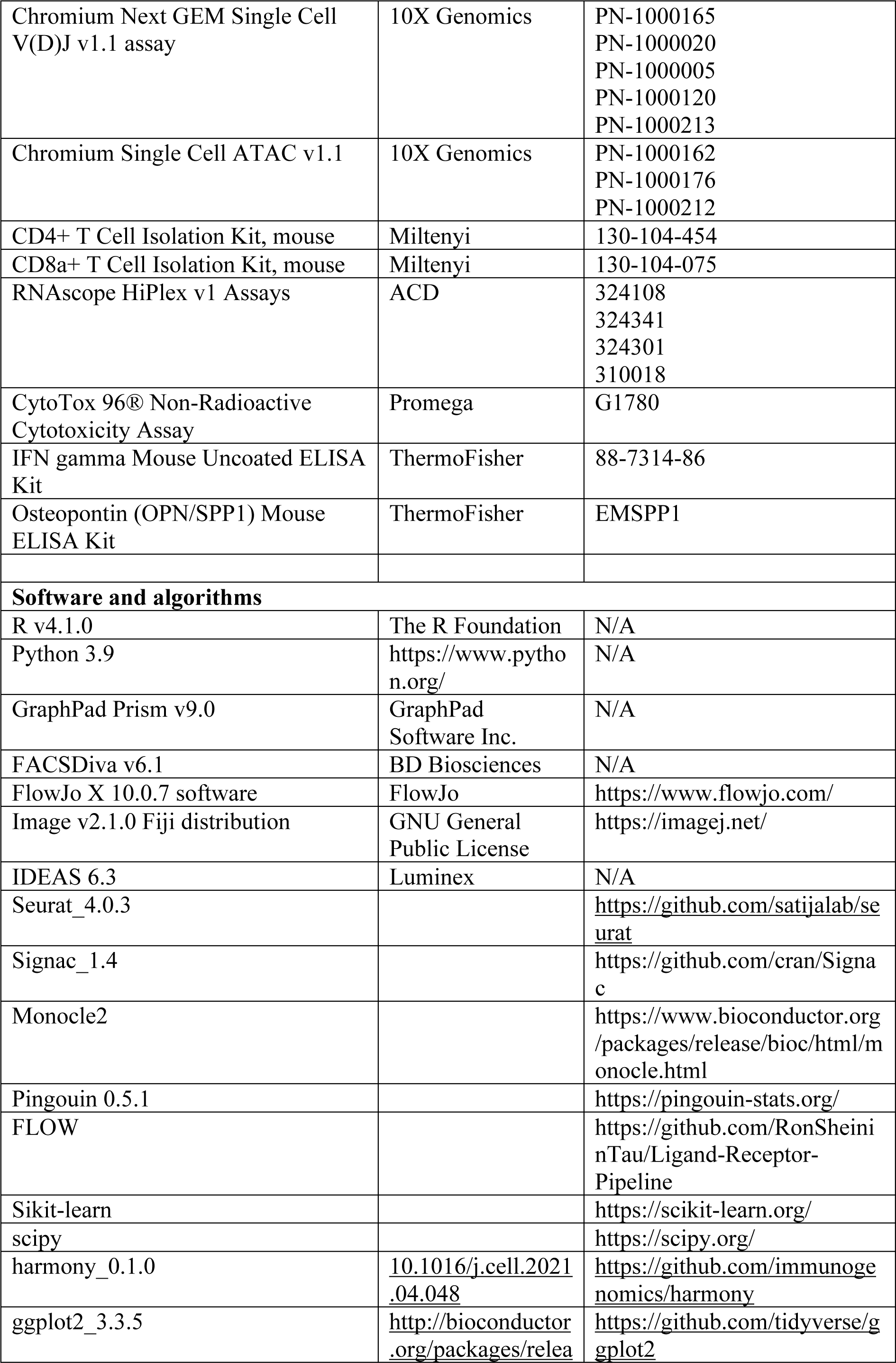
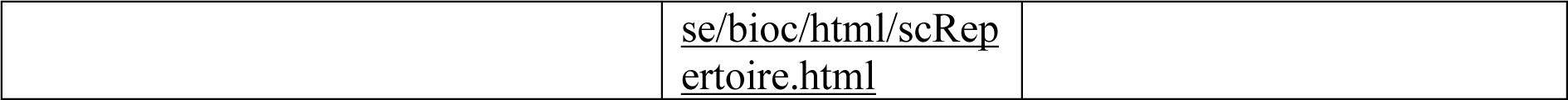

### Resource availability

#### Materials availability

GL261^SIINFEKL^ and GL216^OVA^ cell lines will be made available upon request.

#### Data and code availability

Murine single cell data will be deposited and are publicly available as of the date of publication.

Human single cell data cell data are protected and are not broadly available due to privacy laws. Raw data may be requested from Michael Platten with appropriate institutional approvals.

All original code will be deposited and will be publicly available as of the date of publication.

Any additional information required to reanalyze the data reported in this paper is available from the lead contact upon request.

### Experimental model and subject details

#### Cell lines

Gl261 cells were purchased from the National Cancer Institute. Cell lines were cultured in Dulbecco’s modified Eagle’s medium (DMEM) supplemented with 10% fetal bovine serum (FBS, Sigma Aldrich), 1% penicillin and streptomycin (P/S), at 37 °C, 5% CO2. Cells were routinely tested for mycoplasma contamination. GL261^SIINFEKL^ and GL216^OVA^ cell lines were generated by transfecting GL261 cells with the pMXS-OVA-IRES-blasticidine or pMXS-SIINFEKL-IRES-blasticidine and cultured in DMEM supplemented with 10% FBS, P/S. Transfected cell lines were selected with 9 µg/ml blasticidine (Gibco).

#### Mice

MHCII^flox/flox^ mice were purchased from the Jackson Laboratory and bred at the DKFZ animal facility. Cx3Cr1^CreERT2^ mice were kindly provided by Steffen Jung, Department of Immunology, The Weizmann Institute of Science, Rehovot, Israel. Cx3Cr1^CreERT2^-MHCII^flox/flox^ mice were generated by crossing both strains to achieve homozygous MHCII^flox/flox^ and heterozygous Cx3Cr1^CreERT2^ mice that were used for breeding. For experiments, Cx3Cr1^CreERT2^-MHCII^flox/flox^ and MHCII^flox/flox^ littermates were used. Rosa^CreERT2^ mice were provided DKFZ internally and Rosa^CreERT2^-MHCII^flox/flox^ mice we generated similarly. OT-I and OT-II mice were bred at the DKFZ animal facility.

Mice were housed under Specific and Opportunistic Pathogen Free (SOPF) conditions and under 12-hour day/night cycle. All animal procedures followed the institutional laboratory animal research guidelines and were approved by the governmental authorities (Regional Administrative Authority Karlsruhe, Germany). 8–12-week old mice were assigned to age-matched and sex-matched experimental groups.

#### Human Samples

Human tumor samples were obtained from GBM patients. Written informed consent was obtained by all patients prior to this study conformed to the principles set out in the WMA Declaration of Helsinki and in the Department of Health and Human Services Belmont Report.

Ethical approval for the isolation of brain tumor tissue and single-cell immune analysis was obtained from the Mannheim Medical Faculty Ethics Committee (Reference number 2019-643N).

### Method Details

#### Treatment of mice

Cx3Cr1^CreERT2^-MHCII^flox/flox^ mice or littermates received tamoxifen via oral gavage. For targeting all myeloid cells, 4 days before tumor inoculation, mice were treated with 5 mg tamoxifen (Sigma Aldrich) dissolved in 100 µl corn oil/ethanol (90%/10%, Sigma Aldrich) for 4 consecutive days. Tamoxifen application was repeated every 4-5 days. For a microglia-specific KO, mice were treated with tamoxifen on 5 consecutive days 5-6 weeks before tumor inoculation as described previously (Goldmann et al., 2013).

Immune checkpoint blockade was composed of 250 µg αPD-1 (RMP1-14), 100 µg αCTLA-4 (9D9) and 200 µg αPD-L1 (10F.9G2) (all Bioxcell) blocking antibodies and were intraperitoneally injected starting from day 13 every 3 days for 4 doses. Isotype controls were injected at equal amounts (InVivoMAb rat IgG2a isotype control 2A3, InVivoMAb mouse IgG2b isotype control MPC-11, InVivoMAb rat IgG2b isotype control LTF-2, all Bioxcell). 100 µg agonistic αCD40 antibody (FGK4.5, Bioxcell) was intraperitoneally injected at day 0 and every 4 days until end of the experiment. Isotype control was injected at equal amounts (InVivoMAb rat IgG2a isotype control, 2A3). For CD4^+^ T cell depletion, 250 µg αCD40 (YTS191.1, isotype: LTF-2, both Bioxcell) were intraperitoneally injected 2 days before surgery and repeated every 3-4 days. FTY720 (Cayman) was intraperitoneally injected in PBS at day 15 after surgery when tumors were established. Mice received 1mg/kg FTY or PBS every 2 days.

#### Isolation and culturing of primary murine cells

Spleen and lymph nodes from mice sacrificed by cervical dislocation were mashed through a 70 µm strainer. Erythrocytes were lysed using ACK lysis buffer (Thermo). CD4^+^ from OT-II mice or CD8^+^ T cells from OT-I mice were magnetically separated using MACS negative isolation kits (Miltenyi). In brief, splenocytes were incubated with biotin-labeled antibodies to label unwanted populations. Anti-biotin magnetic beads were added and cells were separated through a column in the magnetic field of a MACS separator (Miltenyi). Splenocytes or isolated T cells were cultured in DMEM, 10% FCS, P/S, 0.1% β-mercaptoethanol (Sigma Aldrich), 1% sodium-pyruvate (Sigma Aldrich), 5 mM HEPES (Invitrogen), Non-essential amino acids (NEAA, Gibco) (T cell medium, TCM).

For bone marrow-derived macrophages, bone and hip from Rosa^CreERT2^-MHCII^flox/flox^ or MHCII^flox/flox^ mice sacrificed by cervical dislocation were isolated and grinded using a mortar in IMDM medium (Thermo Fischer). Cells were mashed through a 40 µm strainer and contaminating erythrocytes were lysed using ACK lysis buffer. For the generation of BMDMs, cells were cultured in IMDM, supplemented with 10% FBS and P/S containing 20 ng/ml M-CSF (Peprotech) and 1 µg/ml 4-Hydroxytamoxifen (Sigma) at 37°C and 5% CO2. 50% of Medium was changed after 72 h and 120 h including M-CSF and 4-Hydroxytamoxifen. Cells were used for assays after 7 days.

#### Intracranial tumor cell inoculation

10^5^ GL261 or GL261^Ova^ or GL261^SIINFEKL^ tumor cells were resuspended in 2 µl PBS and stereotactically implanted into the right hemisphere of 7–14-week-old mice with following coordinates. 2 mm right lateral of the bregma and 1 mm anterior to the coronal suture with an injection depth of 3 mm below the dural surface. A 10 µl Hamilton micro-syringe driven by a fine step stereotactic device (Stoelting) was used. The surgery was performed under anesthesia (Ketamin, 100 mg/kg i.p. und Xylazin, 10 mg/kg i.p) and mice received analgesics for 3 days post-surgery. Mice were checked daily for tumor-related symptoms and mice were sacrificed when stop criteria were met or mice showed signs of neurological deficit.

#### Isolation of tumor infiltrating immune cells

Mice were anesthetized and perfused with PBS. Brains of tumor-bearing mice were extracted and the tumor bearing hemisphere was cut into pieces and digested with 50 µg/ml liberase (Sigma) at 37°C for 30 minutes and subsequently mashed through a 100 µm and 70 µm strainer (Miltenyi). Myelin was removed using a 30% continuous Percoll (GE Healthcare) gradient and centrifugation at 2700 rpm.

#### Flow cytometry for analysis of tumor infiltrating and peripheral immune cells

Murine cells were blocked with rat anti-mouse CD16/32 (0.5 mg per well, eBioscience) and stained with respective antibodies. eFluor 780 fixable viability dye (eBioscience) was used according to manufacturer’s protocol to exclude dead cells. For intracellular staining of cytokines, cells were incubated at 37 °C with 5 mg/mL Brefeldin A (Sigma) for 4 to 6 hours. Intracellular staining was performed using eBioscience Intracellular Fixation & Permeabilization Buffer Set for cytokines (Thermo) or eBioscience™ Foxp3/Transcription Factor Staining Buffer Set for Foxp3 staining (Thermo) according to manufacturer’s protocol. Non-Fixed samples were acquired immediately, and fixed samples were acquired within 48 hours on a FACS Canto II or a BD LSRFortessa (both BD Biosciences).

#### MRI and response criteria

Magnetic resonance imaging (MRI) was carried out at the small animal imaging core facility at DKFZ using a Bruker BioSpec 3Tesla (Ettlingen, Germany) with ParaVision software 360 V1.1. Mice were anesthetized with 3.5% sevoflurane in air. For lesion detection, T2 weighted imaging were performed using a T2_TurboRARE sequence: TE = 48 ms, TR = 3350 ms, FOV 20x20 mm, slice thickness 1,0mm, averages = 3, Scan Time 3m21s, echo spacing 12 ms, rare factor 8, slices 20, image size 192x192. Tumor volume was determined by manual segmentation using Bruker ParaVision software 6.0.1. Response to ICI was determined by using the ratio between tumor volume at 3^rd^ MRI to 2^nd^ MRI. Partial response (PR): ratio < 1, stable disease (SD) 1 < ratio < 1.5, progressive disease (PD) ratio > 1.5.

#### Fluorescence-activated cell sorting of murine T cells for single cell RNA-seq and single cell ATAC-seq

Immune cells were isolated as described above. Murine cells were blocked with rat anti-mouse CD16/32 (0.5 μg per well, eBioscience). Dextramer staining was performed according to manufacturer’s protocol (Immudex): after blocking, cells were incubated with 1:10 dextramer (PE and APC) dilution in 50 µl PBS for 10 minutes. Respective antibodies in PBS were added in 50 µl total volume and stained for 30 minutes. Following antibodies were used: CD45-BV510, CD3-FITC, MHCII-AF700, CD11b-PEDazzle, CD8-PercpCy5.5. eFluor 780 fixable viability dye (eBioscience) was used according to manufacturer’s protocol to exclude dead cells. For scRNA-Seq, cells were pre-incubated for 10 minutes with titrated amounts of TotalSeq C hashing antibodies. Cells were sorted on a BDAria II and BD Aria Fusion through an 85 µM nozzle and 4-way purity. Cells were sorted in 20 µl 0.04% BSA in PBS and kept on ice until processing.

#### Fluorescence-activated cell sorting of tumor infiltrating murine T cells and myeloid cells for co-culture assays

Tumor infiltrating immune cells were isolated as described above. Murine cells were blocked with rat anti-mouse CD16/32 (0.5 μg per well, eBioscience). Cells were stained with respective antibodies in 100 µl PBS. eFluor 780 fixable viability dye (eBioscience) was used according to manufacturer’s protocol to exclude dead cells. Cells were sorted on a BDAria II and BD Aria Fusion through an 85 µM nozzle and 4-way purity in TCM.

For ex vivo killing assays, CD8^+^ T cells were counted and incubated with pre-seeded CFSE labeled GL261^SIINFEKL^ cells for 16 h. Supernatant was subsequently removed and GL261^SIINFEKL^ cells were detached using Accutase (Sigma). Cells were stained with eFluor 780 fixable viability dye (eBioscience) and acquired on a FACS CantoII and quantified using 123count eBeads™ Counting Beads (Thermo).

For ex vivo activation assay, sorted tumor associated bbm and microglia were incubated with OT-II T cells that have been activated for 24 h with 1 µg/ml ISQ peptide and were rested for 3 days. Cells were incubated with 5 µg/ml brefeldinA (Sigma) in TCM for 5 h and intracellularly stained for IFN-γ expression using eBioscience™ Intracellular Fixation & Permeabilization Buffer Set (Thermo). Samples were acquired on a BD FACS Canto II.

#### Cytotoxicity assay of CD8^+^ OT-I T cells against GL261^SIINFEKL^ tumor cells.

Isolated OT-I T cells were cocultured with GL261^SIINFEKL^ or GL261 cells. After 12-36 hours of co-culture an LDH maximum release control of target cells was generated by incubation with 10X Lysis solution (Promega) according to manufacturer’s instructions. Supernatant was used and incubated with CytoTox 96 Reagent (Promega) for 30 minutes in the dark at room temperature. Absorbance at 490 nm was recorded. Minimum LDH release of T cells and Target cells and media background signal were measured for subsequent analysis. Cytotoxicity was calculated as follows: Cytotoxcitiy = (Experimental-effector spontaneous-target spontaneous)/(target maximum-target spontaneous).

#### Isolation and fluorescence-activated cell sorting of human immune cells for single cell RNA-seq

Freshly resected brain tumor tissue was obtained from the university hospital in Mannheim and the university hospital in Heidelberg. Tissue was transported on ice in X-Vivo15 media (Lonza) or PBS (Sigma-Aldrich) and processed within less than three hours after resection. Tumor single cell suspensions were generated by dissecting the tissue into small pieces (2 x 2 x 2 mm) and by subsequently gently mashing the tumor through a 100 µm cell strainer. After filtration through a 70 µm cell strainer, myelin was removed using myelin removal beads II (Miltenyi) and LS columns (Miltenyi) according to the manufacturer’s protocol. Single cell suspensions were cryopreserved in aliquots using 70% FBS (Sigma-Aldrich), 20% X-Vivo20 (Lonza) and 10% DMSO (Sigma-Aldrich) as freezing medium.

For FACS staining, cryovials were thawed, transferred into 10 mL pre-warmed RPMI containing 50 IU/mL benzonase (speed BioSystems), and spun down (350 x g, 10 min, RT). Cell numbers in PBS were counted using trypan blue and turk’s solution before staining for FACS sorting. In short, cells were stained for 10 min at RT with fixable viability dye (AF700, eBioscience). Fc receptors were blocked for 10 min using 10% human serum (Sigma-Aldrich). Staining of cell surface markers was performed as described before. Following antibodies were used: CD235a-APC (clone HI264, Biolegend), CD11b-BB515 (clone ICRF44, BD), CD45-APC-H7 (clone 2D1, BD), and CD3-BV510 (clone HIT3A, BD). After staining, cells were washed, resuspended in FACS buffer (PBS containing 0.5% BSA (Sigma-Aldrich) and 2 mM EDTA (Genaxxon Bioscience)) and transported on ice to a FACS Aria Fusion (BD). Populations of interest were sorted according to the gating scheme into 0.04% BSA in PBS using a 100 µm nozzle (Figure S7A). After FACS sorting, cells were spun down and resuspended in 0.04% BSA in PBS. 12,000 cells were loaded for 10X 5’ single cell sequencing according to the protocols provided by 10X Genomics.

#### Nuclei Isolation and single cell ATAC-Seq preparation

Nuclei isolation was performed using the 10x Nuclei Isolation for Single Cell ATAC Sequencing protocol for low cell input. In brief, dextramer sorted CD8^+^ T cells were pooled for each genotype from individual mice with equal cell numbers. Cells were resuspended in chilled lysis buffer (10 mM Tris-HCl, 10 mM NaCl, 3 mM MgCl2, 0.1% Tween-20, 0.1% Nonidet P40 Substitute (Roche), 0.01% Digitonin (Sigma), 1% BSA (Miltenyi) in nuclease free water) and incubated on ice for 3 minutes. Cells were washed, resuspended in Nuclei Buffer (10x) and 6.000-10.000 nuclei were used for subsequent single cell ATAC library preparation according to the 10x genomics ChromiumNext GEM Single Cell ATAC v1.1 user guide.

#### Co-culture of OT-I, OT-II and BMDMs

CD8^+^ T cells were isolated from OT-I mice and CD4^+^ T cells from OT-II mice using the CD8 T cell or CD4^+^ T cells isolation kit, respectively. BMDMs were generated as described above. 400.00 CD4^+^ and CD8^+^ T cells and 200.000 BMDMs were co-cultured in the presence of 1 µg/ml SIINFEKL and 1 µg/ml ISQ peptide and respective reagents for 3 days. For some experiments, cells were incubated for 7 days and fresh medium was added every 2 days. Cells were stained for intracellular expression of TOX or CD8^+^ T cells were isolated from the co-culture using CD8^+^ T cells isolation kit and rested for 3 days. CD8^+^ T cells were subsequently co-cultured with GL261^SIINFEKL^ cells with 5 µg/ml BrefeldinA (Sigma) and stained for intracellular cytokine production. Target cell killing was assessed with LDH assay (Promega) after 24 h of co-culture as described above.

#### OT-I activation assays

OT-I splenocytes were isolated as described above and stimulated with 1 µg/ml SIINFEKL peptide and 50 iU IL-2/ml. 1 µg/ml OPN or 1 µg/ml αCD29 were added. 50% of the medium was exchanged every 2 days supplemented with IL-2, OPN and αCD29. TOX and NFAT2 expression was analyzed using flow cytometry after 3 and 7 days. For NFAT2 translocation, cells were intracellularly stained for NFAT2 as described above using eBioscience™ Foxp3/Transcription Factor Staining Buffer Set for Foxp3 staining (Thermo). Samples were acquired on a ImageStream X MKII (Luminex) equipped with following lasers: 375 nm, 405 nm, 488 nm, 561 nm, 582 nm, 642 nm, 785 nm. 5000 events/sample were acquired. Compensation was performed using single stains and IDEAS 6.3 analysis software (Luminex). Cells were gated for best focus, single cells and CD8-FITC and DAPI positivity. Nuclear localization of NFAT2 was assessed using the Nuclear Localization feature implemented in the IDEAS software.

#### Immunofluorescence

Mice were anesthetized and perfused with PBS. Brains were isolated and quickly frozen in OCT tissue Tek (Sakura) and stored at -80°C. 8 µM sections were collected on SuperFrost slides (Epredia) using a cryotome (Leika). A hydrophobic barrier was drawn around the sample and slides were incubated in -20°C cold methanol for 10 minutes. Samples were subsequently blocked for 1 h with 10% normal goat serum (NGS, Cell Signaling Technology) in 0,1 % Tween+ PBS at room temperature. Slides were incubated with following antibodies at 4°C over night: human anti-CD3 (1:300, GeneTex), rat anti-mouse CD8a (2,5 µg/ml ThermoFischer) and rat anti-mouse MHCII (1:200, Thermo). Primary antibodies were incubated overnight at 4°C. Samples were washed three times with TBS-T and incubated with respective secondary antibodies (all 1:300) for 1 hour at room temperature: Alexa Fluor 488 labeled goat anti-rat, Alexa Fluor 633 labeled goat anti-rat, Alexa Fluor 633 labeled goat anti-rabbit, (all Invitrogen). For subsequent staining with fluorochrome conjugated antibodies slides were washed as previously described and then incubated with FITC-conjugated rat anti-mouse F4/80 (BM8, BioLegend) at a dilution of 2,5 µg/ml for 2 hours at 4°C. Slides were mounted using Fluoromount-G with DAPI (Invitrogen) and let dry for 2 h. Images were acquired on a microscopy Zeiss LSM 700 or an Olympus VS200 slide scanner with 40x magnification. Images were processed and analyzed with Fiji software. Quantification was performed by automated counting of DAPI positive nuclei in signal area overlays above a predefined threshold using Fiji software or by manual counting.

#### In situ hybridization in human GBM tissue

Freshly resected brain tumor tissue was obtained from the university hospital in Mannheim and directly frozen in liquid nitrogen. 10 µm sections were prepared as described above. In situ hybridization of RNA-probes was performed using RNAscope^TM^ HiPlex Assay (ACD) according to manufacturer’s protocol. Briefly, slides were fixed with 4% PFA for 60 min and dehydrated with 50%, 70% and 100% ethanol. A hydrophobic barrier was drawn around the sample and slides were incubated with protease IV for 30 min. Subsequently, probes were hybridized according to manufacturer’s protocol. Slides were counterstained with ProLong Gold Antifade mountant (ThermoFisher) and imaged the next day using an Olympus VS200 slide scanner with 40x magnification. For the next imaging rounds, fluorophores were cleaved and new fluorophores were hybridized for 4 imaging rounds in total and. Fluorophores for three RNA-probes were imaged each round (12 in total). The probes are listed in supplementary table 1. For image registration the HiPlex image registration software (ACD) was used. Images were processed and analyzed with Fiji software. Regions of interest were defined by tissue matched integrity with approximately 150 x 150 µm in size. MHCII expression was quantified using the mean intensity of the respective channel. Mean intensity was used as threshold for MHCII^high^ and MHCII^low^ regions. Quantification was performed by manual counting of cells positive for the respective probe.

#### Adoptive transfer of CD4^+^ OT-II T cells

CD4^+^ T cells were isolated from OT-II mice as described above. 2x10^6^ cells in 150 µl PBS were injected into the tail vein of GL261-Ova bearing mice at day 20 after tumor inoculation. After 10 days, mice were anesthetized and spleen and lymph nodes were extracted. Mice were perfused with 1x PBS and the tumor bearing hemisphere was collected. Cells were isolated as described above and stained with respective antibodies.

#### Data processing, clustering and annotation murine scRNAseq

The data was pre-processed, normalized, clustered and annotated using Seurat R package for single cell analysis as was previously described at (Hao et, al 2021). Briefly, the pipeline consists of the following steps. **LogNormalize**: each feature counts for each cell are divided by the total counts for that cell and multiplied by a scale factor. **Dimensionality reduction:** PCA and tSNE are calculated from the scale normalized data matrix, where each feature normalized expression is scaled across the cells. The number of PCs for the clustering was manually selected based on elbow plot showing the gain in variance with each additional vector. **Clustering:** First, we calculated the k-nearest neighbors and constructed the KNN graph, in the reduced PCA space. On that graph a modularity score is optimized using the Leiden clustering method (Traag, Waltman, and van Eck 2019). **Cluster annotation:** was done manually by the use of known cell population markers and projection of known cell type gene signatures on the UMAP plots.

#### Differentially expressed genes between clusters

The FindAllMarkers function in Seurat was used to find marker genes that were differentially expressed between clusters. This function identifies differentially expressed genes between two groups of cells using a Wilcoxon Rank Sum test with limit testing chosen to detect genes that display an average of at least 0.25-fold difference (log-scale) between the two groups of cells and genes that are detected in a minimum fraction of 0.25 cells in either of the two populations. This step was intended to speed up the function by not testing genes that are very infrequently expressed.

#### Visualization of single cell data

To generate UMAP plots of single cell profiles, the scores along the 15 significant PCs described above were used as input for the R implementation of UMAP, by the RunUMAP function in Seurat. Heatmaps were generated using DoHeatmap function in Seurat for the top 10 DE genes in each cluster.

#### Single-cell gene signature scoring

Single-cell gene signature scoring was used to identify subset of cells that are associated with a given genes list that represent a biological condition, as described in (Kurtulus et al. 2019). Scores were computed by first sorting the normalized scaled gene expression values for each cell followed by summing up the indices (ranks) of the signature genes. A contour plot which takes into account only those cells that have a signature score above the indicated threshold was added on top of the UMAP space, in order to further emphasize the region of high scoring cells.

#### Signature score compression between different cell populations

Given a signature score value for each cell in a subset, we could qualify the difference of score means between different populations. For the compression of two populations, an unpaid t-test was used, and for three or more population a one-way Anova test was applied with Tukey’s post hoc analysis to detriment the significance between every pair of subsets in the data. Using this statistical compression, we could determine if a given gene signature is enriched in a cell population relative to the others.

#### Trajectory analysis

To obtain the Trajectory graph, and calculate the pseudotime, Monocle2 DDR-Tree was used on the CD8^+^ T cell cluster. The algorithm first applies the Revers Graph embedding method in order project the high dimensional data onto a low dimensional space, then a Minimum

Spanning Tree is calculated on set of cluster centroids to obtain the tree structure, finally algorithm gives each cell a pseudotime value, which represents the distance (relative time) of every cell in the dataset from a group we defined as the root of the tree.

In order to ensure the quality of the result, we calculate the rolling mean signature score over the pseudotime of early versus late CD8^+^ cells. This signature was obtained from independent dataset, of CD8^+^ T cells harvested from tumors at two different timepoints (see Methods). The strong correlation observed, is evidence that that trajectory was indeed able to depict the differentiation of cells over time.

#### Distribution over pseudotime

After obtaining the pseudotime value for each cell in the data, we can calculate the distribution of cell over the pseudotime axis and visualize it using a histogram. This calculation can be done for each separate biological condition; thus, we could learn the difference in differentiation stages between the different conditions.

#### Rolling mean over pseudotime

Given a signature score or a single gene expression value and a pseudotime value for every cell in the data, we first order the cell by their pseudotime value. In order to reduce the noise attributed to sparse gene expression values from single cell data, a rolling mean of the gene expression value or signature score was calculated with a window size of 200 cells. Thus, we could plot the mean score over the pseudotime and view the expression profile over the suggested differentiation routs.

#### Ligand receptor detection

In order to identify ligand-receptor putative interactions between different cell populations using Single Cell RNA seq data we applied the FLOW framework. Briefly the algorithm receives a clustered dataset and for each pair of cell clusters, a signal sender and signal receiver clusters are defined and differentially expressed ligands and receptors are identified. In order estimate the ability of a given receptor to propagate its signal downstream, we calculate the maximum flow from each receptor to a set of transcription factors using a PPI network. Finally, permutation tests are used to estimate the significance of the downstream activation score. Using the FLOW framework, we can also identify how the signal from different receptors can converge onto a given downstream transcription factor. In addition, as we calculate the multi-source maximum flow from the set of receptors to the TF using the PPI network, we can also identify which receptor can affect a specific transcription factor of interest.

#### ATAC-Seq Analysis

The ATAC-seq data was analysis using the Signac and Seurat packages in R. First the data was normalize using the term frequency-inverse document frequency (TF-IDF) method. Then, Singular value decomposition (SVD) was applied on the normalized matrix, and the second to the 30^th^ dimensions was selected in order to represent the data onto a low dimensional space, then UMAP was apply to visualize the data.

In order to find DE peaks between the conditions, we run Logistic regression test using FindMarkers function of Seurat. Finally, we looked at the genes that were in the distance of 10,000 bp from every DE peak we found.

#### Bulk RNA expression analysis and TCGA analysis

Bulk RNA dataset GSE121810 was downloaded from Gene Expression Omnibus dataset and imported into R (v.4.1.2). Count matrix was normalized and log-transformed using DESeq2 (v1.34). The expression of MHC II related genes (HLA and CD74) was summed to represent MHCII expression. The patient cohort was divided by median expression of MHCII in MHCII^high^ and MHCII^low^.

Correlation of genes from the TCGA GBM dataset was performed using the GEPIA2 webtool (Tang et al., 2019).

#### Single cell RNA-seq analysis of human samples

Single cell datasets were aligned using cellranger (v 6.1.5). Count matrices of single cell dataset were then imported into R(v4.1.2) for further analysis. QC was done individually using miQC (v1.2) and Seurat (v. 4.0.5) was used to merged and normalize the data. Harmony (v0.1) was then subsequently used to integrate the dataset. Non-T and non-myeloid cells were excluded from the analysis. Canonical markers were then used to identify and annotate the cell type. The MHCII Signature and Exhaustion Signature were calculated using AddModuleScore function in Seurat. The MHCII Signature included following genes: HLA-DRA,HLA-DRB5,HLA-DRB1,HLA-DQA1,HLA-DQB1,HLA-DQB1-AS1,HLA-DQA2,HLA-DQB2,HLA-DOB,HLA-DMB,HLA-DMA,HLA-DOA,HLA-DPA1,HLA-DPB1,CD74. The Exhaustion Signature included following genes: PDCD1, TIGIT, LAG3, CTLA4, TOX, HAVCR2. Multiple correlation was then performed using Hmisc (v4.6) and plotted using corrplot (v0.96).

#### Quantification, statistical analysis and figures

Data are represented as individual values or as mean ± SEM. Applied statistical tests are indicated in each figure legend. Statistics were calculated using GraphPad Prism 9.0 or the R packages Hmisc for multiple correlation.

Some figures were created with biorender.com.

**Figure S1.**
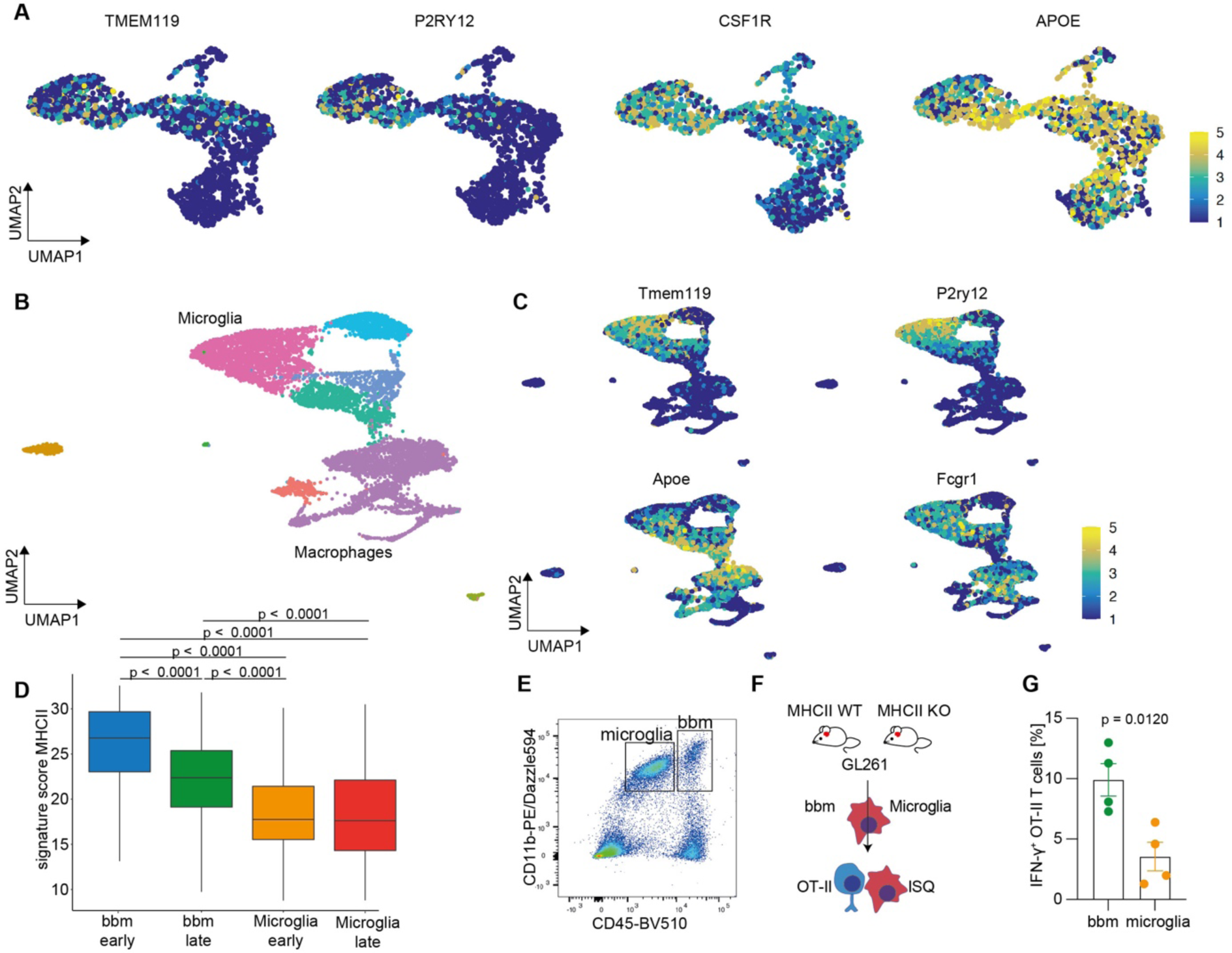
MHC class II expression in human and murine tumor infiltrating immune cells. (A) Feature plots of microglia and bbm associated genes in human scRNA-seq dataset from Figure 1(C). (B) UMAP of murine CD45^+^/CD3^-^ cells sorted from early (day 14) and late (day 26) GL261 tumors. (C) Feature plots of murine microglia and bbm associated genes (D) Quantification of MHCII antigen presentation signature from (B). Statistical significance was tested by one-way Anova test with Tukey’s post hoc analysis. (E) Experimental overview. CD45^high^/CD11b^+^ bbm and CD45^low^/CD11b^+^ microglia were sorted and co-cultured with pre-activated OT-II CD4^+^ T cells. (F) Sorting gates for bbm and microglia. (G) IFN-γ^+^ OT-II cells after co-culture with ISQ peptide loaded bbm and microglia sorted from GL261 tumors. Bars indicate mean ± SEM. Statistical significance was tested by unpaired t test.

**Figure S2.**
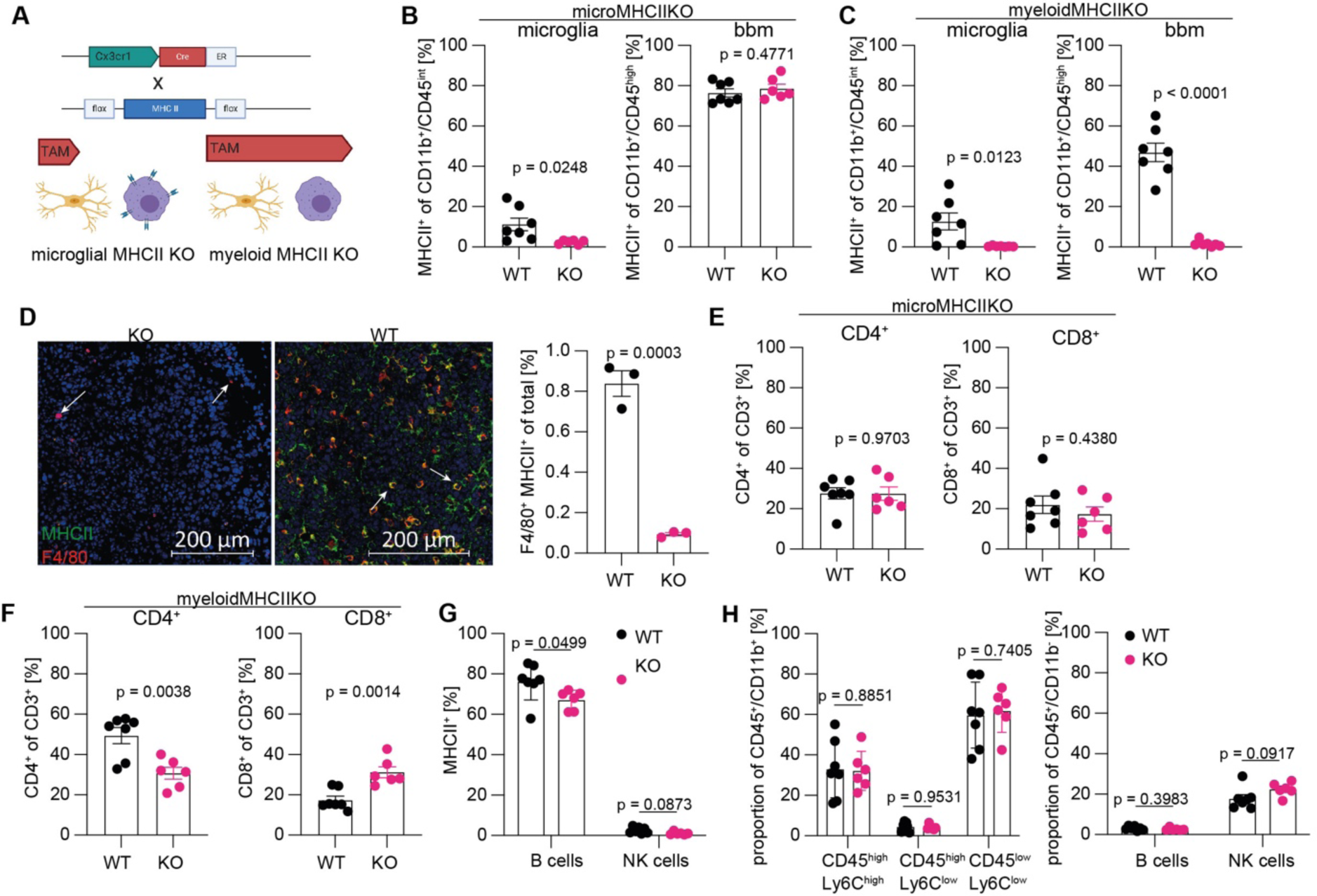
Validation of Cx3cr1^CreERT2^-MHCII^flox/flox^ mice (A) Schematic representation of the Cx3cr1^CreERT2^ inducible MHC class II knockout using timed dosing of tamoxifen. (B) MHC class II expression on microglia and bbm after microMHCIIKO tamoxifen treatment schedule. (C) MHC class II expression on microglia and bbm after myeloidMHCIIKO treatment schedule. (D) Representative immunofluorescence imaging of MHC class II expressing F4/80^+^ myeloid cells (left), quantification (right). (E) Analysis of immune cells infiltrating of GL261 tumors from MHCII WT and MHCII KO mice under myeloidMHCIIKO tamoxifen treatment schedule. (F) Analysis of immune cells infiltrating of GL261 tumors from MHCII WT and MHCII KO mice under microMHCIIKO tamoxifen treatment schedule. (G) MHC class II expression on B cells and NK cells under myeloidMHCIIKO tamoxifen treatment schedule (H) Quantification of CD3^-^ immune cell subsets abundance under myeloidMHCIIKO tamoxifen treatment schedule. (B), (C), (E)-(H) analyzed by flow cytometry. (B)-(H) bars indicate mean ± SEM. Statistical significance was tested by unpaired t test.

**Figure S3.**
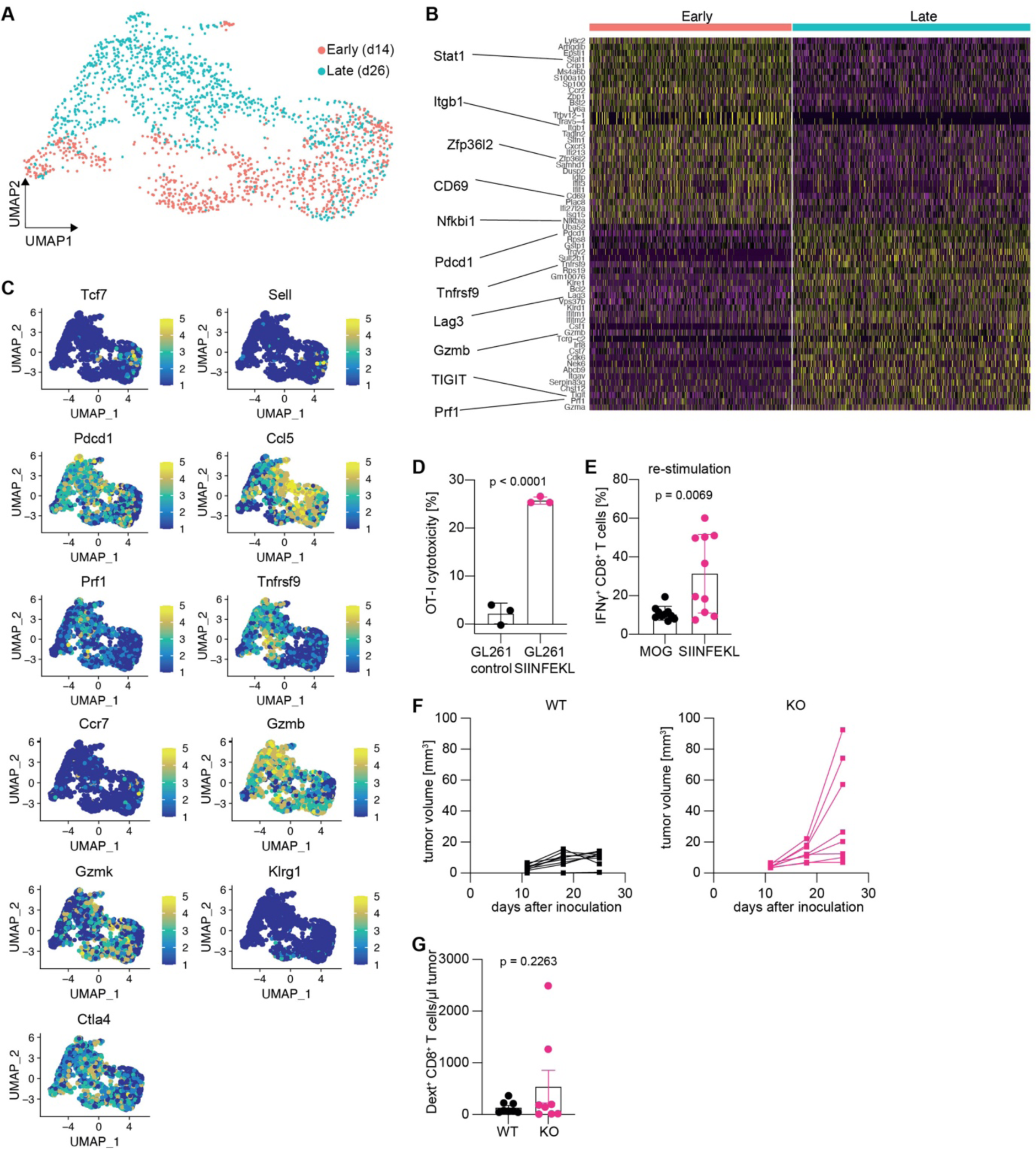
CD8 T cells drive anti-glioma immune response in GL261 WT and GL261^SIINFEKL^ tumors. (A) UMAP of CD8^+^ cells sorted from early (day 14) and late (day 26) GL261 tumors. (B) Heatmap of the top 30 marker genes separating early and late tumors with representative genes annotated on the left. (C) Representative density plots of exhaustion/activation markers in early and late tumors. (D) Cytotoxicity of OT-I T cells against the GL261^SIINFEKL^ cell line, measured by LDH release. (E) IFN-γ production of CD8^+^ T cells isolated from GL261^SIINFEKL^ tumors or cervical lymph nodes restimulated with SIINFEKL peptide. (F) Individual tumor growth curves of GL261^SIINFEKL^ tumors in MHCIIWT and myeloidMHCIIKO mice. (G) Absolute numbers of Dext^+^ CD8^+^ T cells in GL261^SIINFEKL^ tumors. (D), (E), (F) bars indicate mean ± SEM. Statistical significance was tested by unpaired t test.

**Figure S4.**
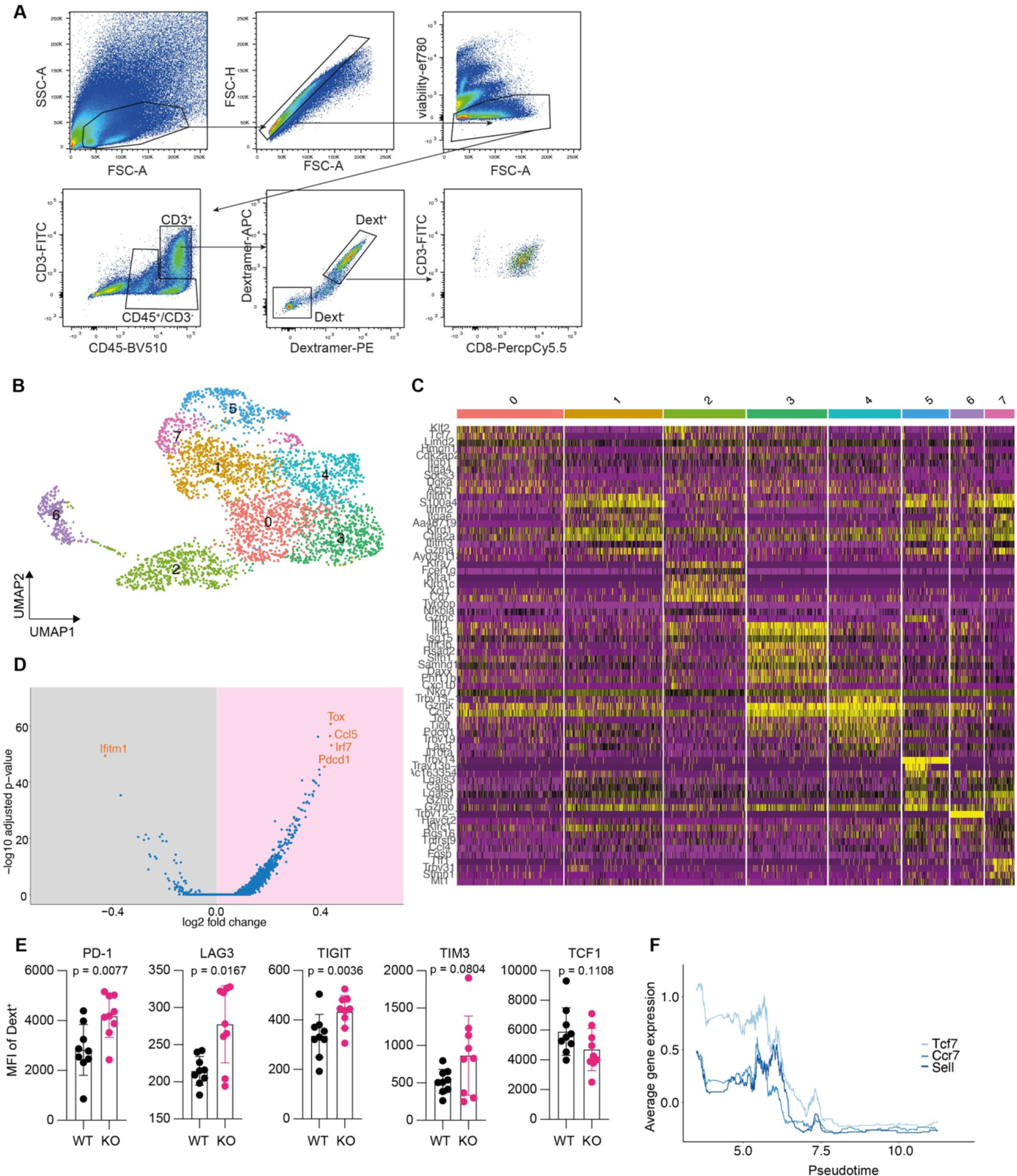
Loss of myeloid MHC class II expression drives tumor reactive T cells into distinct transcriptional dysfunction state. (A) Sorting strategy for murine immune cells from GL261^SIINFEKL^ tumors at day 20 after tumor inoculation. (B) Clustered UMAP representation of CD8^+^ T cells sorted from GL261^SIINFEKL^ tumors. (C) Heatmap of top10 cluster defining genes (D) Volcano plot of differentially expressed genes in CD8^+^ T cells from MHCIIWT and myeloidMHCIIKO mice. (E) Quantification protein expression of selected markers on Dext^+^ CD8^+^ T cells, related to Figure (3 E). Bars indicate mean ± SEM. Statistical significance was tested by unpaired t test. (F) Rolling mean of stem-like markers along the pseudotime trajectory in Dext^+^ CD8^+^ T cells.

**Figure S5:**
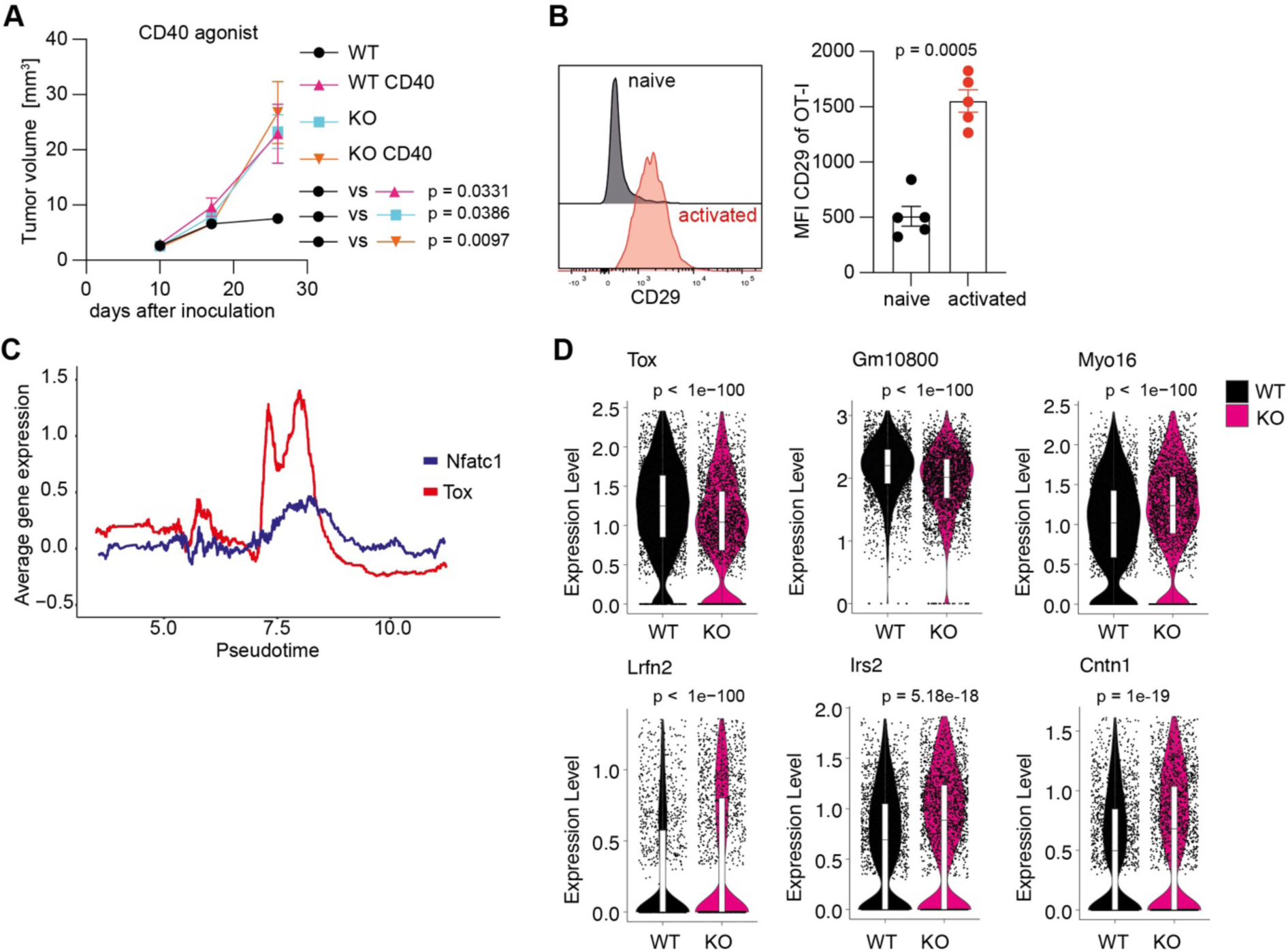
Loss of MHCII leads to Nfatc1-mediated Tox expression. (A) Tumor volume measured by MR imaging after agonistic CD40 antibody treatment in MHCIIWT and myeloidMHCIIKO mice. (B) Representative flow cytometry plot for CD29 protein expression on naïve and activated OT-I T cells (left) and quantification of CD29 MFI (right). (C) Rolling mean of Tox and Nfatc1 expression along the pseudotime trajectory, see also Figure 3J, K. (D) Differentially accessible genes in on Dext^+^ CD8^+^ T cells sorted from GL261^SIINFEKL^ tumors. (A, B) bars indicate mean ± SEM. (A) Statistical significance of tumor volumes was tested by one-way ANOVA with Tukey’s test at day 27. (B), (D) Statistical significance was tested by unpaired t test.

**Figure S6.**
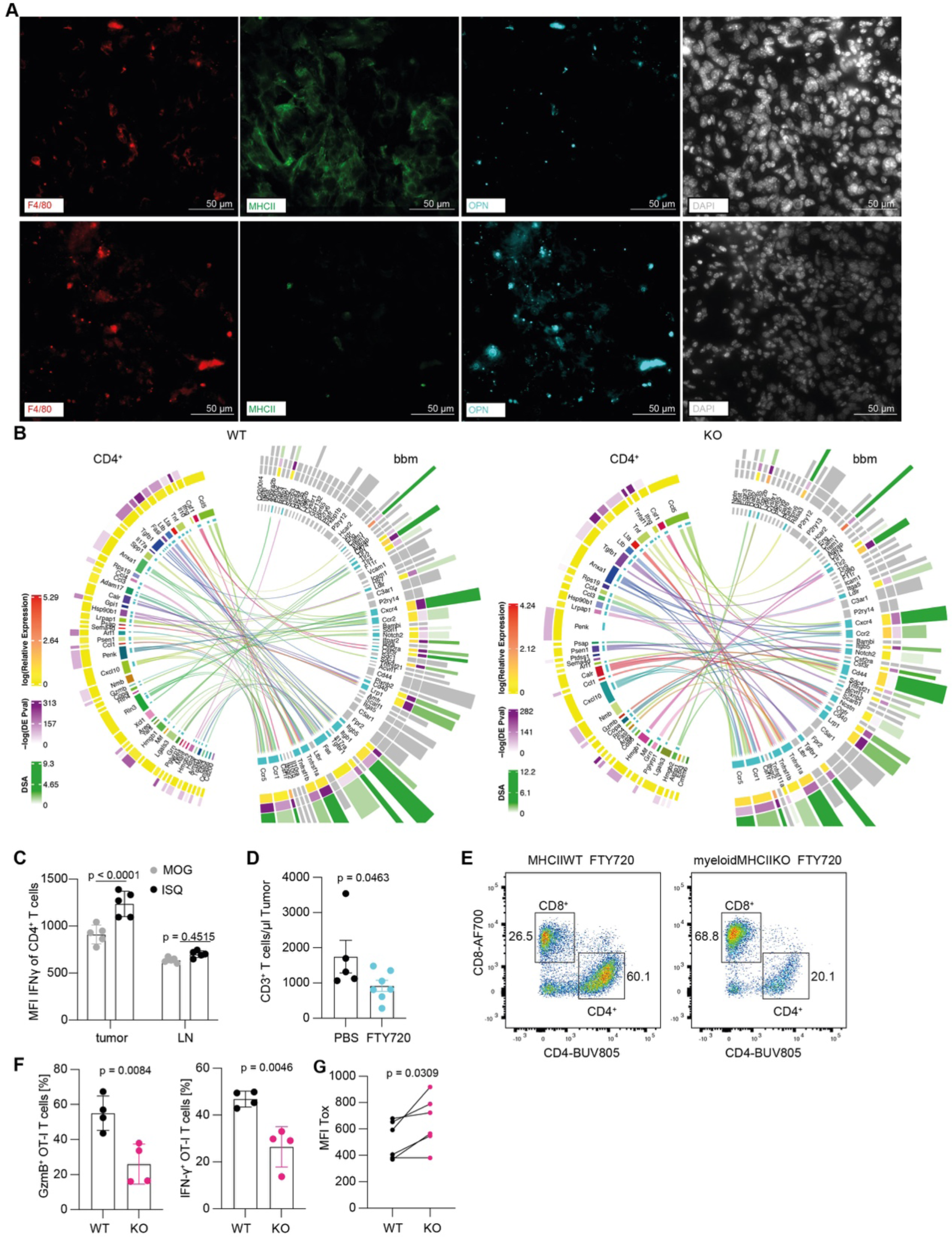
OPN expression is regulated by intratumoral MHCII and CD4^+^ T cells (A) Single channels of representative immunofluorescence images of GL261^SIINFEKL^ tumors. See also Figure 5A. (B) Interaction map of CD4^+^ T cells and bbm from MHCII WT and MHCII KO mice. (C) MFI of IFN-γ in CD4^+^ T cells from GL261^Ova^ tumors or cervical lymph nodes (LN) (D) Absolute numbers of tumor infiltrating T cells in FTY720 or PBS treated mice. (E) Representative flow cytometry plot of CD3^+^ T cells of FTY720 treated MHCIIWT and myeloidMHCIIKO mice. (F) GzmB and IFN-γ expression after recall of OT-I T cells primed in the presence or absence of MHCII and OT-II T cells, see also Figure M. (G) MFI of Tox after 7 days in OT-I T cells primed in the presence or absence of MHCII and OT-II T cells, combined data from 2 individual experiments, see also Figure 5M. (C), (D), (F), (G) bars indicate mean ± SEM. (C), (D), (F) Statistical significance was tested by unpaired t test. (G) Statistical significance was tested by paired t test.

**Figure S7.**
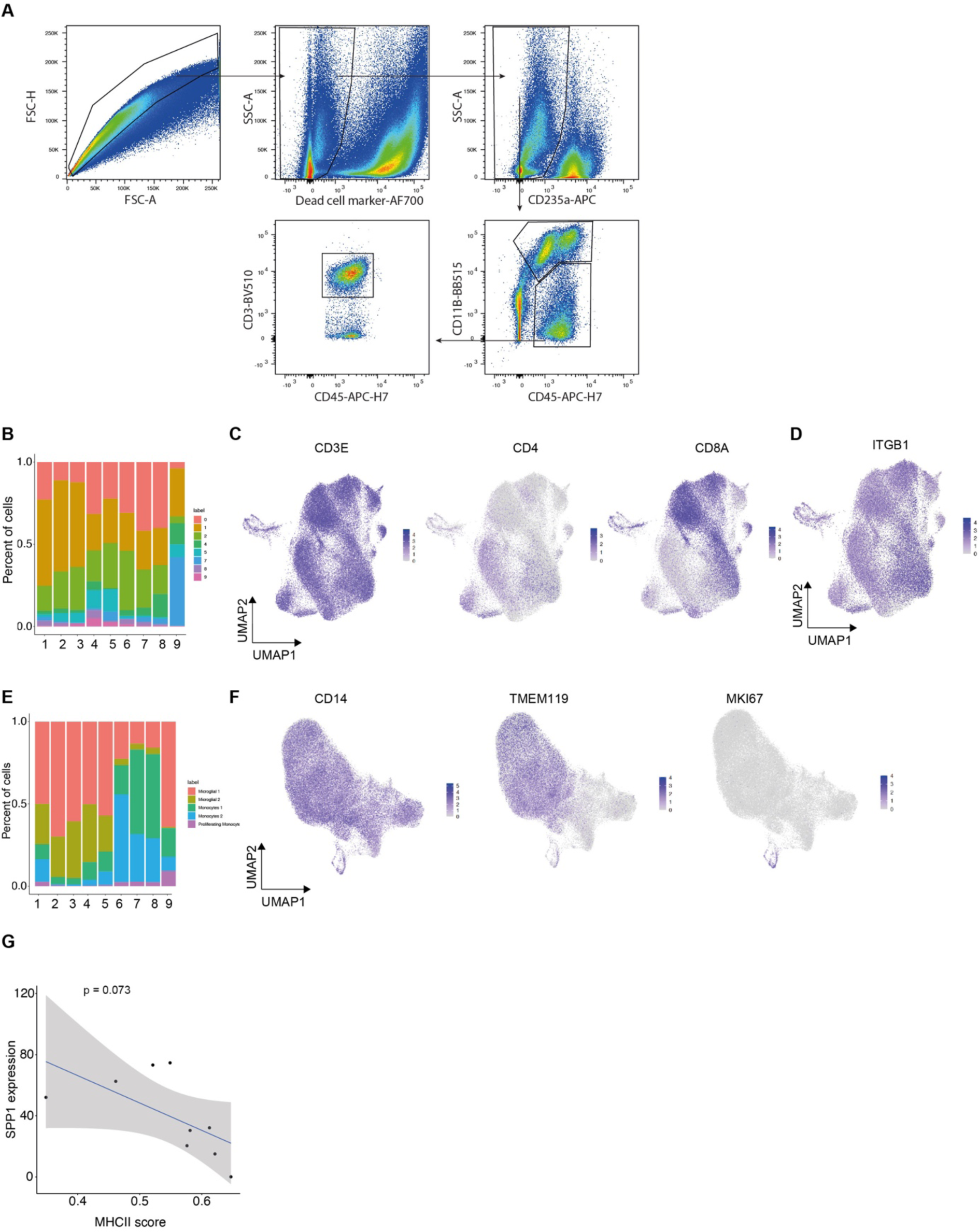
ScRNA-seq of human GBM. (A) Sorting strategy for immune cells from human GBM samples. (B) Visualization of the distribution of T cell cluster among patients. (C) Feature plots for CD3e, CD4 and CD8 RNA expression. (D) ITGB1 expression on T cells from (A). (E) Visualization of the distribution of myeloid cluster among patients. (F) Feature plots for CD14, TMEM119 and Mki67 RNA expression. (G) Correlation of SPP1 expression and MCHII score in MHCII^high^ bbm per patient.

